# Engineering an Anaerobic Microenvironment to Empower Hydrogenase-Catalyzed Hydrogen Therapy for Diabetic Wound Healing

**DOI:** 10.1101/2025.12.01.691506

**Authors:** Haishuo Ji, Yaling Wang, Kexin Yao, Junjie Li, Hang Luo, Wangzhe Li, Yanxin Gao, Wenjin Li, Qi Xiao, Tin Pou Lai, Chunxiao Chen, Xueying Li, Qian Peng, Chunqiu Zhang, Baofa Sun, Liyun Zhang

**Author notes:** Corresponding authors. *E-mail addresses* (Q. Peng), (C. Zhang), (B. Sun), (L. Zhang).

## Abstract

The inherent oxygen sensitivity of hydrogenases has long restricted their applications to anaerobic systems. Here, we present a hybrid peptide–nanocluster hydrogel that establishes a self-sustaining anaerobic microenvironment to enable hydrogenase-catalyzed hydrogen therapy under aerobic conditions. This system integrates a custom-designed Fmoc-KYF peptide hydrogel with integrated silver nanoclusters (AgNCs), which host an engineered [NiFe]-hydrogenase (3S1C-Hyd-2). Mechanistic studies combining molecular dynamics simulations and spectroscopic analysis revealed that hydrophobic pockets within the peptide network trap oxygen molecules, while photoexcited AgNCs rapidly deplete invasive O_2_. Our design ensures stable hydrogen evolution, effectively overcoming the oxygen sensitivity that has long restricted hydrogenase applications in biomedical contexts. In vitro, the in-situ-generated hydrogen effectively scavenges reactive oxygen species, suppresses pro-inflammatory cytokine production. In a diabetic mouse model, this light-activated system markedly accelerates wound closure (87% by day 11), reduces oxidative stress, promotes macrophage polarization toward a reparative phenotype, enhances angiogenesis, and upregulates tissue repair pathways, as confirmed by transcriptomic profiling. Our work establishes a new bioengineering paradigm for constructing anaerobic microenvironments through synergistic material–enzyme–light interactions, offering a versatile platform for hydrogenase-enabled therapeutic interventions in oxidative stress–related diseases.

## 1. Introduction

Hydrogenases are a diverse family of metalloenzymes that catalyze the reversible conversion between protons and molecular hydrogen (2H⁺ + 2e⁻ ↔ H_2_), thereby playing a central role in microbial energy metabolism [1]. These enzymes are widely distributed across bacteria [2], archaea [3], and some eukaryotes [4], where they occur either in soluble form or associated with membranes, typically localized in the periplasm, cytoplasm, or specialized organelles [5]. By mediating hydrogen oxidation and evolution, hydrogenases enable organisms to harness H_2_ as an energy source or to dispose of excess reducing equivalents, thereby maintaining intracellular redox homeostasis. Depending on their subcellular localization and physiological context, they are functionally adapted for hydrogen uptake, hydrogen production, or the generation of transmembrane proton gradients that drive cellular bioenergetics. Notably, hydrogenases exhibit exceptional catalytic performance, with turnover frequencies approaching 10⁴ s⁻¹ and minimal overpotentials comparable to platinum catalysts, despite relying on earth-abundant nickel and/or iron at their active sites. This unique combination of efficiency, selectivity, and sustainability has spurred intense interest in their mechanistic study, biotechnological exploitation, and the design of biomimetic hydrogen-evolving systems [6]. However, a critical limitation of hydrogenases is their sensitivity to oxygen (O_2_) [7]. The widespread application of hydrogenases under aerobic conditions remains a significant challenge.

Strategies to enhance the O_2_ tolerance of hydrogenases can be broadly divided into two main categories: bioengineering and material-assisted approaches [8]. Bioengineering methods employ genetic modification techniques to improve the intrinsic resistance of hydrogenases to oxygen inactivation [9,10]. In contrast, combining enzymes and synthetic materials creates an anaerobic microenvironment with electron donors and physical barriers using electronic media and functional nanostructures, protecting hydrogenase from oxygen exposure [11–16]. However, these strategies rely on toxic electron mediators, such as methyl viologen (MV), which limit their applicability in biological contexts beyond energy conversion systems.

[NiFe]-hydrogenases are a diverse and intriguing group of hydrogenases characterized by a heterodimeric structure consisting of a large subunit containing the NiFe active site and a small subunit harboring one to three iron-sulfur (FeS) clusters that facilitate electron transfer between the active site and the protein surface [17]. A representative structure of *Escherichia coli* [NiFe] hydrogenase 2 (Hyd-2) exhibits remarkable capacity for integration into functional materials for light-driven H_2_ evolution systems [8,18–24]. However, Hyd-2 is highly oxygen-sensitive in photo-H_2_ production systems, even trace amounts of O_2_ leading to rapid deactivation [19]. This renders the Hyd-2 catalytic system challenging to apply in biomedical fields characterized by the widespread presence of oxygen. Herein, we report a novel strategy to protect Hyd-2 in a therapeutic dressing from O_2_-induced inactivation by employing a custom-engineered hybrid peptide hydrogel, as illustrated in Scheme 1. In this system, silver nanoclusters (AgNCs), integrated into the peptide backbone and conjugated to Hyd-2, serve as multifunctional components that act simultaneously as redox mediators, light harvesters, and molecular linkers. Upon photoexcitation, the AgNCs function as efficient electron donors, facilitating electron transfer through the FeS cluster relay to promote H_2_ production. This protective hydrogel matrix safeguards the catalytically active NiFe site against oxygen exposure via a “Three-Tier Protection, Dual-Mechanism” approach that combines a physical diffusion barrier with a self-activated catalytic O_2_-scavenging process. The resulting Hyd-2-based photocatalytic system demonstrates substantial therapeutic potential, as evidenced by anti-inflammatory effects at the cellular level and accelerated wound healing in diabetic mice. Our findings have significant implications for advancing innovative hydrogenase-based biomedical applications beyond the energy sector.

**Scheme 1.**
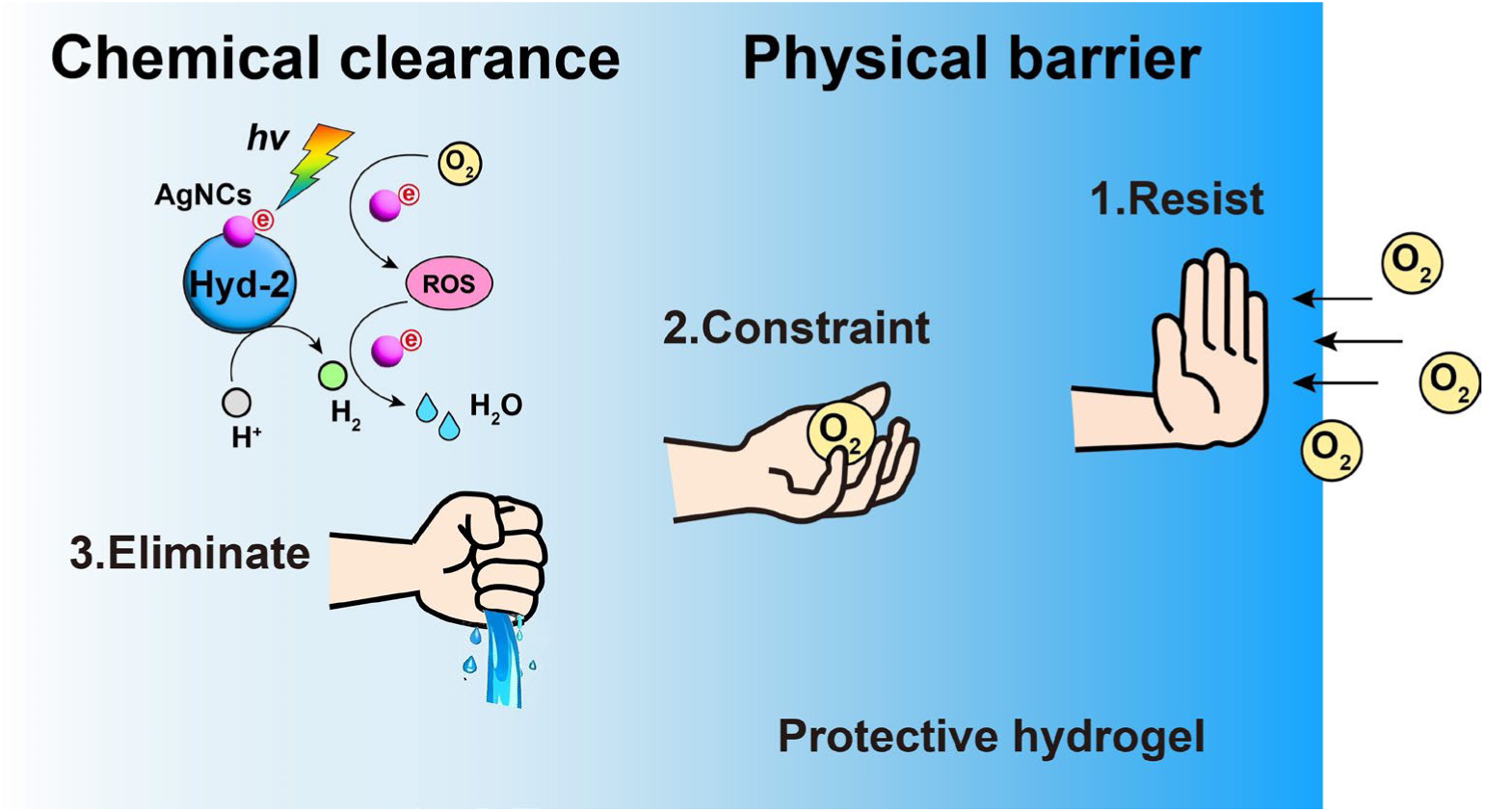
Schematic illustration of the engineered anaerobic microenvironment that enables hydrogenases to sustain high catalytic activity under aerobic conditions.

## 2. Results and Discussion

### 2.1 Design of a Protective Hydrogel Matrix

To effectively protect Hyd-2 from oxygen exposure, we developed a hydrogel matrix composed of a self-assembled peptide for the encapsulation of Hyd-2 (Fig. 1a). Drawing inspiration from the well-characterized Fmoc-diphenylalanine (Fmoc-FF) peptide hydrogelator, which forms nanofibers and generates hydrogels in aqueous solutions through π–π interactions between aromatic rings [25–27], a *de novo* hydrogelator with a distinct structural configuration was designed. Considering the high hydrophobicity and strong aggregation tendency of Fmoc-FF in aqueous environments, which necessitates the use of organic solvents to dissolve the peptides at high concentrations—potentially detrimental to enzyme activity—we hypothesized that replacing one phenylalanine with tyrosine (Y) to form YF, thereby incorporating a hydrophilic group (–OH), would modulate the thermodynamic process of self-assembly and promote hydrogel formation similar to that of FF [27]. Additionally, to facilitate electrostatic interactions between the peptides and electronegative AgNCs, lysine (K), which carries a positive charge under physiological conditions, was incorporated into the peptide sequence, yielding KYF. Accordingly, the tripeptide Fmoc-KYF was synthesized via solid-phase peptide synthesis, and its structure is illustrated in Fig. 1a. The purity and structure of Fmoc-KYF were validated using HPLC, NMR, and mass spectrometry (Fig. S1-S4). Notably, Fmoc-KYF exhibited moderate solubility and exceptional stability in aqueous environments.

**Fig. 1.**
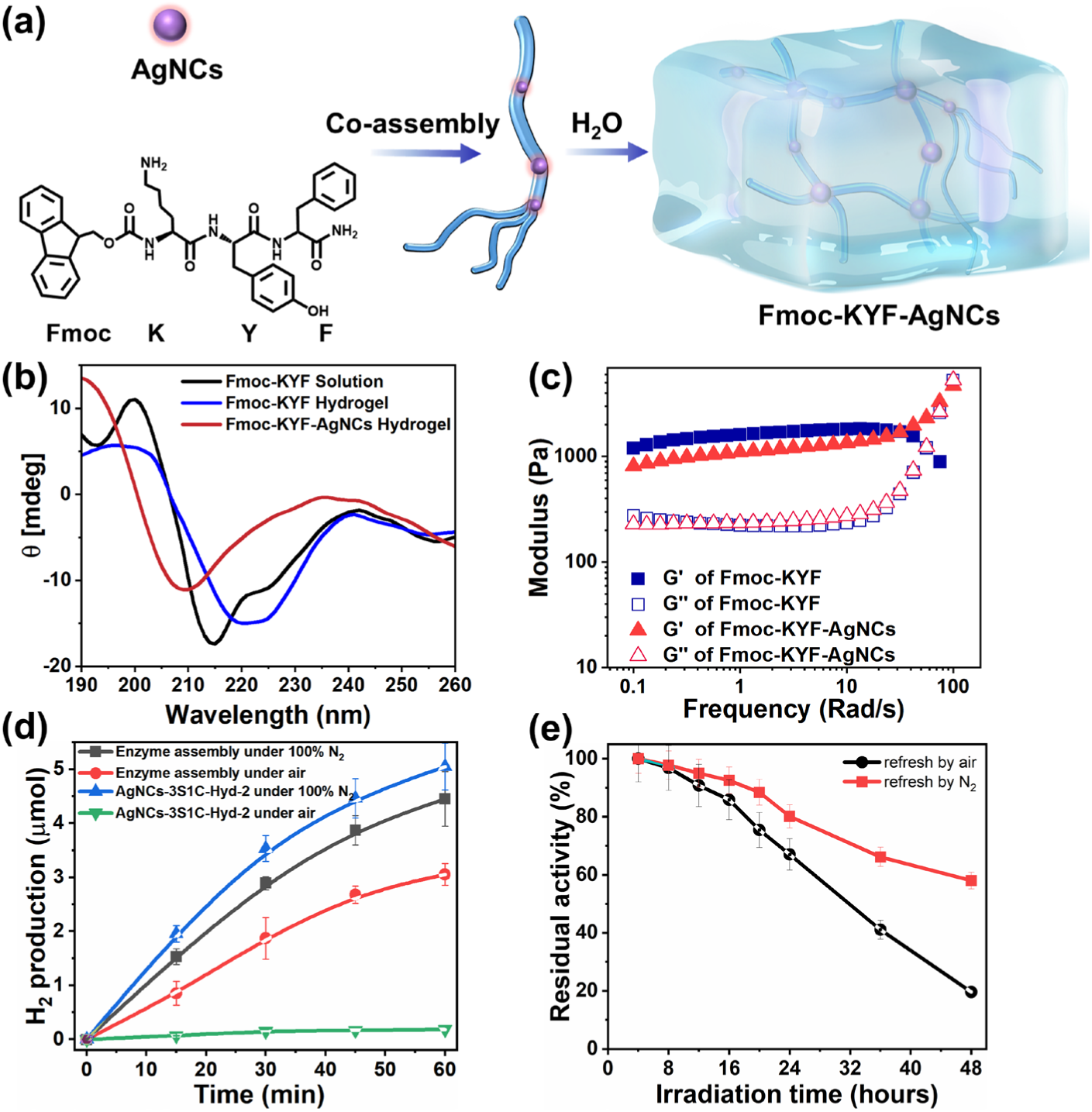
(a) Schematic of co-assembly between Fmoc-KYF peptide and silver nanoclusters (AgNCs) under mild conditions, yielding a self-supporting hydrogel network. (b) Circular dichroism spectra of Fmoc-KYF alone (black for the solution and blue for the NaCl induced hydrogel) and after incorporation of AgNCs (red) to the hybrid hydrogel. (c) Rheological analysis of the hydrogel (5 mg mL⁻¹ peptide, 0.5 mg mL⁻¹ AgNCs). (d) Time-dependent H_2_ production under aerobic (air) vs. anaerobic (N_2_) conditions for Hyd-2 assembly (black/red) and AgNCs-3S1C-Hyd-2 (blue/green). (e) Operational stability over 48 h continuous irradiation with periodic headspace renewal. 2 mL total volume containing 0.25 nmol 3S1C-Hyd-2, 5 mg mL⁻¹ Fmoc KYF, 0.5 mg mL⁻¹ AgNCs, 0.1 M TEOA, pH 7.0.

Circular dichroism (CD) spectroscopy of the Fmoc-KYF solution revealed a peak at 215 nm, indicating a β-sheet conformation (Fig. 1b, black curve) [27]. As anticipated, Fmoc-KYF was capable of forming a peptide hydrogel directly in response to salt without the need for any organic solvent (Fig. S5), as evidenced by the minimum at 220 nm, which suggested a transition to a super α-helical arrangement (n → π* transitions, Fig. 1b, blue curve) [25]. Upon the addition of AgNCs(Fig. S6 and S7), the CD spectrum exhibited maxima at 190 nm and minima at 208 nm, characteristic of α-helical π → π* transitions (Fig. 1b, red curve) [25,28]. Subsequently, rheological analysis of the hydrogels showed that the storage modulus (G′, ∼1100 Pa) was approximately five times greater than the loss modulus (G″, ∼230 Pa), confirming the formation of a robust yet soft hydrogel (Fig. 1c) [29]. Notably, the incorporation of AgNCs into the hydrogel did not result in substantial alterations to its mechanical properties, thereby preserving the favorable attributes of the hydrogel material for use as a therapeutic dressing [30,31].

### 2.2 Proof of Concept for 3SC1-Hyd-2-Catalyzed H_2_ Production under Aerobic Conditions

To verify the feasibility of the designed anaerobic microenvironment construction strategy, we first evaluated the photocatalytic hydrogen production performance following hydrogenase assembly. In our previous work, we engineered a Hyd-2 quadruple variant (C433S, C432S, C197S, Y222C; hereafter 3S1C-Hyd-2, Fig. S8) by introducing a surface-exposed cysteine (Fig. S9, C222) near the distal [4Fe-4S] cluster for AgNCs linkage, enabling direct electron transfer and enhanced light-driven H_2_ evolution [21]. H_2_ evolution was confirmed by bubble formation (Fig. S10) and then quantified via gas chromatography (Fig. 1d), with an initial rate of 4.45 μmol h⁻¹ under anaerobic conditions. Under aerobic conditions, the rate decreased by ∼68% to 3.05 μmol h⁻¹ (black trace), whereas without the hydrogel, the assembly lost over 95% activity within 15 minutes in the presence of air (green trace). Comparative analysis reveals that hydrogen production (the hydrogen content detectable by gas chromatography) under aerobic conditions is reduced compared to anaerobic conditions, which may be attributed to the following factors: (1) partial inactivation of hydrogenases caused by oxygen; (2) preservation of enzymatic activity with concurrent consumption of a portion of the generated hydrogen, leading to reduced gas chromatography detection signals. Systematic variation of enzyme concentration, pH, and electron donors (Fig. S11-S13) underscored the hydrogel matrix’s crucial role in optimizing H_2_ production. Under optimized conditions (2 mL total volume containing 0.25 nmol 3S1C-Hyd-2, 5 mg mL⁻¹ Fmoc KYF, 0.5 mg mL⁻¹ AgNCs, 0.1 M TEOA, pH 7.0), as shown in Fig. 1e, continuous irradiation over a 48-hour period, with headspace flushing performed every 4 hours, resulted in an I₅₀ (activity reduced by 50%) value of 32 hours under aerobic conditions. These findings also indicate that during the initial phase of the reaction (as shown in Fig. 1d), the observed decrease in hydrogen production under aerobic conditions is not attributable to enzyme inactivation.

### 2.3 Mechanistic Analysis of Anaerobic Microenvironment Formation

The photocatalytic hydrogen production performance under ambient conditions, as experimentally demonstrated above, confirms the feasibility of the designed system. Subsequent studies will focus on elucidating the underlying mechanism of this design strategy. We have known that during peptide self-assembly, the balance of intermolecular interactions among different molecules, including hydrogen bonding, electrostatic interactions, and aromatic-aromatic interactions, influences nanostructure formation and leads to distinct morphologies [32]. First, transmission electron microscopy (TEM) was used to reveal the unique microstructural architecture. Fig. 2a and 2b showed a well-defined fibrous network in the Fmoc-KYF and Fmoc-KYF-AgNCs hydrogel, with fibrils ranging from 10 to 50 nm in diameter. Each nanofiber comprised multiple intertwined peptide chains. AgNCs aggregated into nanoparticles (5–10 nm in diameter) distributed along the peptide nanofibers. *In situ* liquid-phase atomic force microscopy (AFM) imaging further demonstrated a multi-stage assembly process. Initially, Fmoc-KYF molecules formed primary assembly nanofibers, which further twisted into large bundles (approximately 30 nm in diameter, Fig. 2c, d). As illustrated in Scheme 1, all these distinctive structures constitute the fundamental basis for the matrix’s capacity to achieve physical oxygen barrier properties and suppress deep diffusion.

**Fig. 2.**
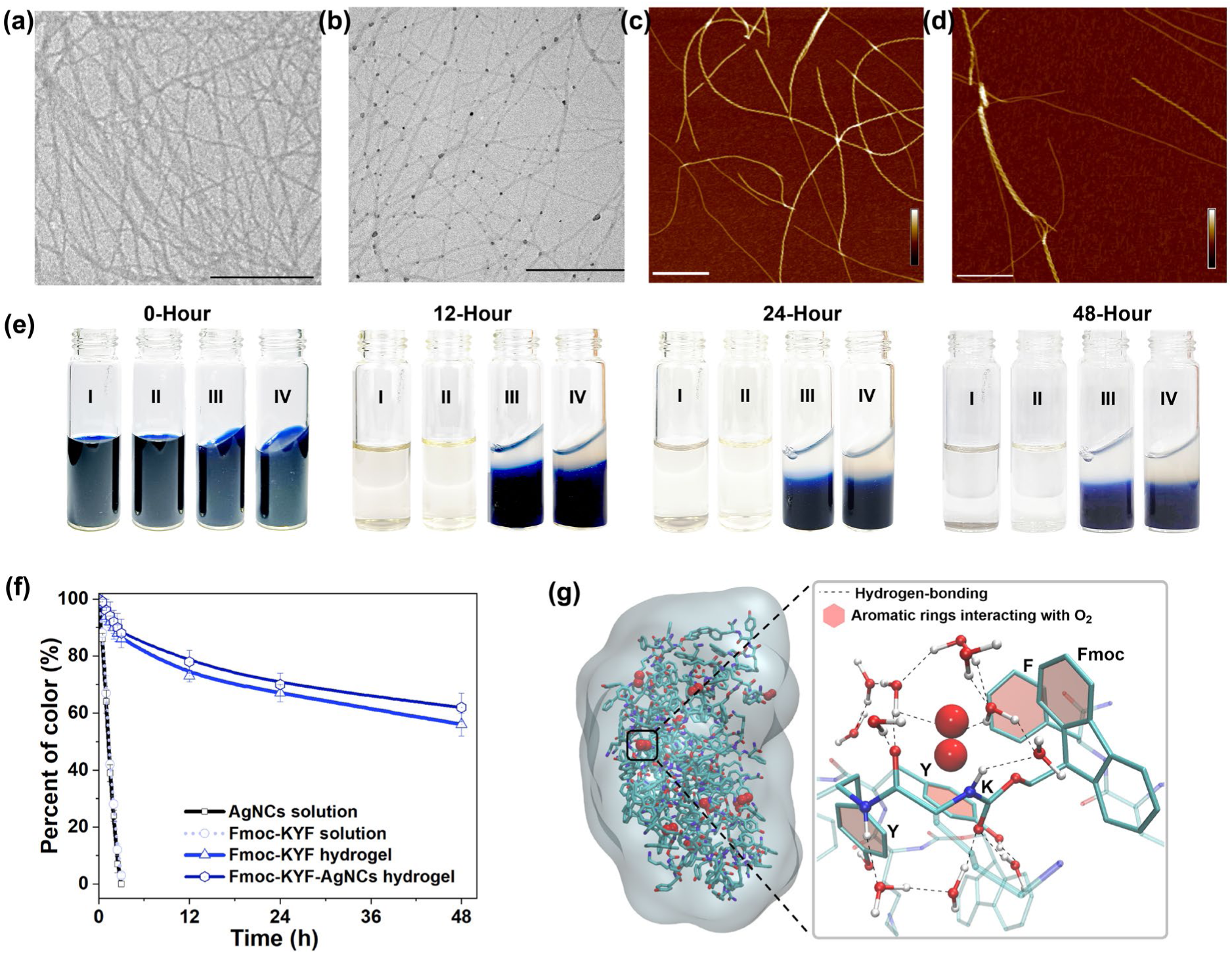
The TEM images of the Fmoc-KYF hydrogel (a) and Fmoc-KYF-AgNCs hydrogel (b), where AgNCs appear as dark clusters. Scale bar: 500 nm. AFM topography images showing fibril persistence length for the Fmoc-KYF hydrogel (c) and Fmoc-KYF-AgNCs hydrogel (d). Scale bar: 200 nm. (e) Time-resolved MV⁺· decolorization under ambient O_2_. (I-IV) 2 mM MV²⁺ in AgNCs solution, Fmoc-KYF solution, Fmoc-KYF hydrogel and Fmoc-KYF-AgNCs hydrogel. (f) Quantitative analysis of color retention kinetics for the MV⁺· radical (λ = 605 nm) in Fmoc-KYF solution or hydrogel. Data were normalized to the initial time point (t = 0) and are presented as mean ± SD (n = 3). (g) Molecular dynamics simulation of O_2_ diffusion constraints (red spheres) within the Fmoc-KYF matrix.

Subsequently, the methyl violet fading assay (MV^+·^ +O_2_ → MV^2+^) was employed to evaluate the physical barrier effect of Fmoc-KYF-AgNCs against oxygen permeation. MV^+·^ (violet color, absorption peak at 605 nm) was generated by reducing 2 mM MV^2+^ (colorless) with sodium dithionite, producing a bright violet color at 0 h (Fig. 2e). The color of MV^+·^ solution completely faded within 2 hours upon exposure to air oxidation (Fig. 2e, vials I and II). In contrast, both NaCl-induced Fmoc-KYF and hybrid Fmoc-KYF-AgNCs hydrogels retained 60% of the MV^+·^ color intensity after 48 hours (Fig. 2e, vials III and IV), which indicates that the three-dimensional nanostructure of the hydrogels significantly restricts oxygen diffusion (Fig. 2f). Notably, among all tested concentrations, the 5 mg mL⁻¹ gels provided significantly better O_2_ protection for MV^+·^ (Fig. S14). We hypothesize that at higher concentrations, Fmoc-KYF fibril aggregation occludes O_2_ binding sites, thereby reducing the effective surface area.

To further elucidate the O_2_–hydrogel interaction, we performed molecular dynamics simulations of Fmoc-KYF hydrogels in explicit water with O_2_ and H_2_ using Gromacs 2018.8 [33] and the OPLS-AA force field (Fig. S15-S18) [34]. The simulated hydrogel structure demonstrated that O_2_ was effectively encapsulated by the fibrils, whereas H_2_ exhibited negligible binding (Fig. S17, Supporting Videos V1 and V2). Analysis of the simulated binding conformations revealed a plausible O_2_ binding mode (Fig. 2g and S18). O_2_ interacts with one Fmoc group, two tyrosine rings, one phenylalanine ring, and multiple water molecules. These hydrophobic pockets feature a unique hydrogen-bond network between these water molecules and peptide amide group (from lysine residue) that tightly confines O_2_ molecules, thereby greatly impeding their diffusion into the hydrogel interior.

Subsequent experiments were designed to investigate the role of AgNCs in oxygen scavenging. Initially, a series of experimental procedures were conducted to demonstrate the stable assembly of AgNCs on polypeptide scaffolds. Photoluminescence titrations, ζ-potential measurements, and FTIR spectroscopy collectively confirmed that AgNCs interact with Fmoc-KYF through electrostatic interactions and/or potential ligand exchange (Fig. S19-S21) [35]. As shown in Fig. 3a, the hybrid hydrogel exhibits superior luminescent performance, providing a catalytic foundation for efficient oxygen removal via chemical pathways. Under illumination, the Fmoc-KYF-AgNCs hydrogel generates significant reactive oxygen species (ROS) in the presence of O_2_, as indicated by the strong fluorescence of the ROS-specific probe 2’,7’-dichlorofluorescin diacetate (DCFDA, red curve in Fig. 3b) [36]. In contrast, no ROS generation was observed in the polypeptide hydrogel alone (blue curve, Fig. 3b). Given that the LUMO and HOMO of AgNCs are −0.55 and 1.25 V vs NHE, respectively [37] and the LUMO is more negative than the oxygen reduction potential for O_2_/·O_2_− (−0.33 V vs NHE, pH 7) and O_2_/H_2_O_2_ (0.28 V vs NHE, pH 7) [38], AgNCs satisfy the thermodynamic requirements for generating ·O_2_^−^ and H_2_O_2_ (Fig. 3c). This result aligns with prior studies on the photocatalytic conversion of O_2_ to ROS by Ag9-AgTPyP [38].

The introduction of 3S1C-Hyd-2 significantly reduced fluorescence intensity (green curve, Fig. 3b), indicating that the produced hydrogen effectively neutralizes ROS generated by AgNCs under light irradiation. Replacing the hydrogenase-based hydrogen production system with a saturated molecular hydrogen solution (∼800 μM) under identical conditions led to a substantial decrease in ROS fluorescence (purple curve, Fig. 3b). This strongly supports that molecular hydrogen enables the photocatalytic reduction of O_2_ to water. These findings also provide a clear explanation for the reduced hydrogen production observed under aerobic conditions (Fig. 1d), indicating that hydrogen contributes to the establishment of an anaerobic microenvironment by scavenging oxygen, thereby protecting hydrogenase activity from oxidative degradation. Although the reaction between hydrogen and oxygen to form water under ignited conditions is known, our work innovatively introduces a “light-triggered ignition” mechanism that enables efficient oxygen scavenging under mild conditions. This design successfully establishes a local microenvironment that maintains high catalytic activity of the hydrogenase, allowing continuous photocatalytic hydrogen production under ambient air or physiological conditions—laying a critical foundation for its biomedical applications.

**Fig. 3.**
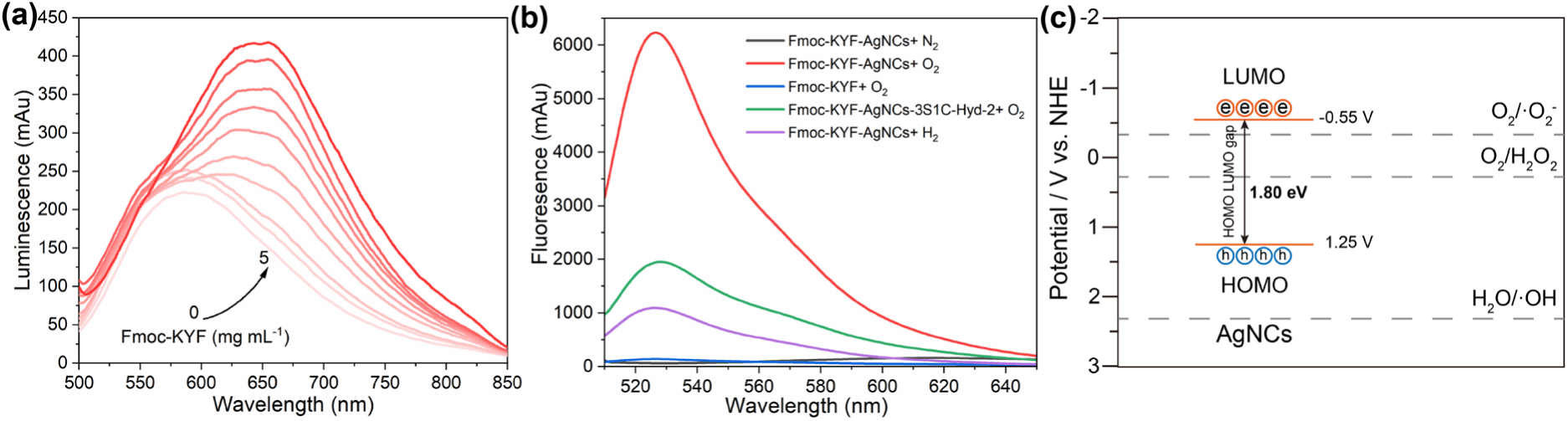
(a) Photoluminescence spectra of AgNCs under Fmoc-KYF peptide titrated. The concentration of AgNCs is 60 μM, concentrations of Fmoc-KYF are 0-5mg mL^-1^. The excitation wavelength is 460 nm. (b) ROS generation profiles (as indicated by DCFDA fluorescence, λₑₓ: 488 nm) under illumination for 10 minutes under various conditions. (c) The band positions of AgNCs with respect to the ROS formation potential.

### 2.4 ROS scavenging and anti-inflammation

H_2_ has garnered increasing attention as a therapeutic agent for inflammatory conditions due to its ability to selectively neutralize cytotoxic species such as hydroxyl radicals and peroxynitrite [39], thereby mitigating ROS-induced cellular damage at its source [40]. Inspired by the aforementioned results demonstrating that 3S1C-Hyd-2 can achieve efficient light-driven hydrogen production under aerobic conditions, we subsequently evaluated the physiological effects of hydrogen generated by 3S1C-Hyd-2 using a lipopolysaccharide (LPS)-induced inflammatory model. LPS stimulation in RAW264.7 macrophages is known to elevate intracellular ROS (•OH, ONOO⁻, H_2_O_2_) and proinflammatory cytokines (e.g., IL-1β, IL-6) [41], leading to a robust inflammatory response. Cells were treated with either Fmoc-KYF-AgNCs hydrogel or 3S1C-Hyd-2 assembly under light irradiation, alongside concurrent LPS exposure for 12 hours; untreated cells and LPS-stimulated cells served as controls. Confocal fluorescence imaging using DCFDA (a probe for ROS) and DAPI (a nuclear stain) revealed a substantial increase in green fluorescence in LPS-treated cells, indicating excessive accumulation of intracellular ROS (Fig. 4a). Furthermore, quantitative DCFDA assays corroborated these observations (Fig. S22). Notably, treatment with the 3S1C-Hyd-2 assembly resulted in a significantly greater reduction in intracellular ROS compared to the Fmoc-KYF-AgNCs hydrogel alone, which can be attributed to enhanced in situ hydrogen generation.

**Fig. 4.**
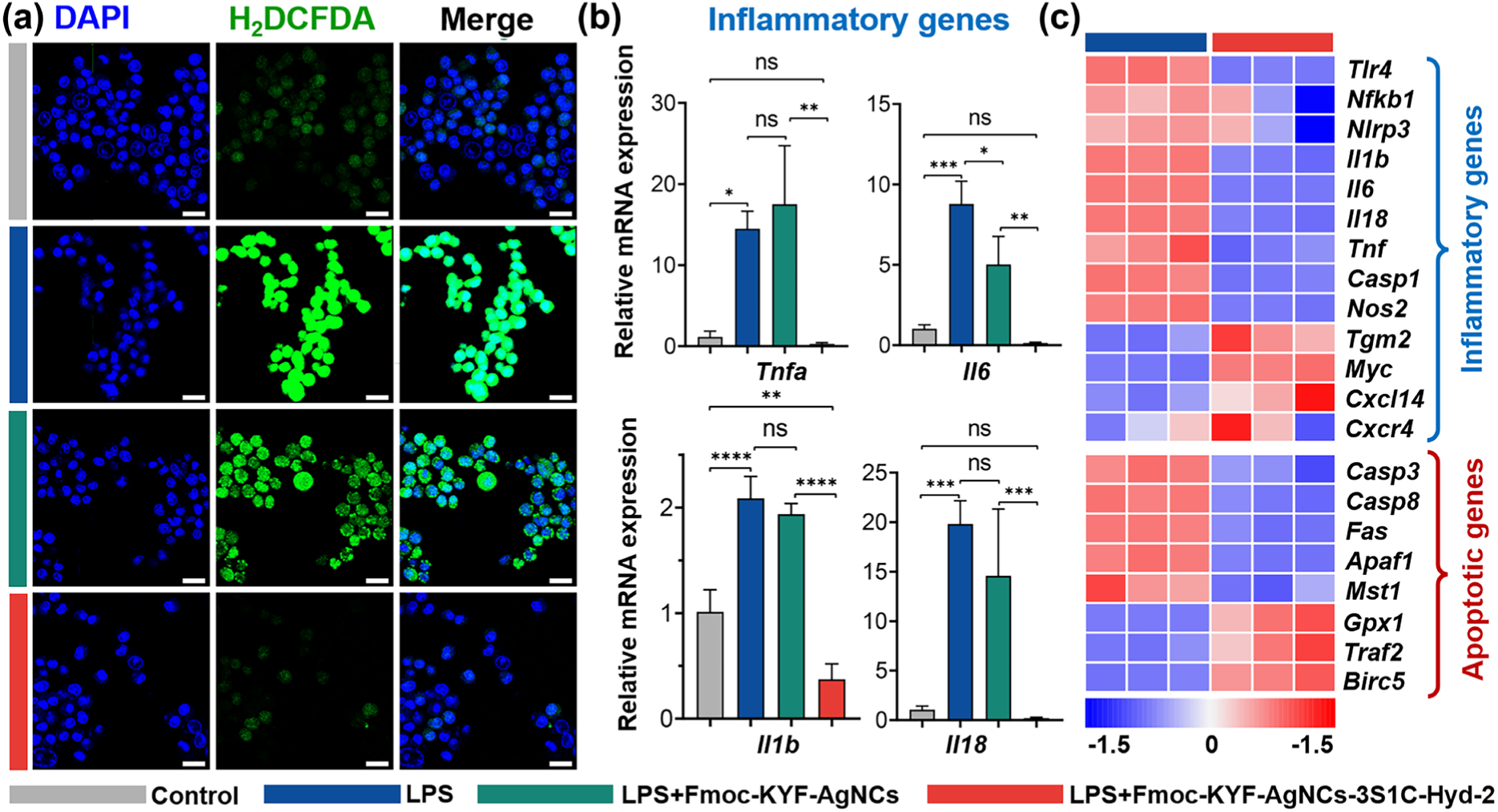
ROS scavenging and anti-inflammatory effects of Hyd-2 assembly. (a) Confocal fluorescence images showing intracellular ROS levels in LPS-stimulated RAW264.7 macrophages following treatment with different hydrogels. scale bar: 20μm. (b) Real-time quantitative PCR analysis of representative inflammatory gene expression. Statistical significance: ns, no significance, *P<0.05, **P<0.01, ***P<0.001, ****P<0.0001. (c) Heatmap showing the relative expression of representative inflammatory and apoptotic genes in cells treated by Hyd-2 assembly from RNA-seq data (n=3).

Gene expression analysis by real-time quantitative PCR demonstrated that 3S1C-Hyd-2 assembly significantly suppressed mRNA levels of *Tnfa, Il1b, Il18,* and *Il6*, relative to Fmoc-KYF-AgNCs hydrogel (Fig. 4b), indicating effective inhibition of LPS-induced proinflammatory cytokine production. Concurrently, mRNA levels of anti-apoptotic marker (*Bcl2*) were elevated (Fig. S23), aligning with enhanced cell viability observed in CCK-8 assays (Fig. S24). Bulk RNA sequencing further revealed that in LPS-stimulated macrophages, treatment with 3S1C-Hyd-2 assembly downregulated key pro-inflammatory (*Tlr4, Nfkb1, Nlrp3, Il1b, Il6, Il18*) and pro-apoptotic genes (*Casp3, Casp8, Fas, Apaf1, Mst1*), while upregulating anti-inflammatory (*Tgm2, Myc, Cxcl14, Cscr4*) and anti-apoptotic genes (*Gpx1, Traf2, Birc5*) (Fig. 4c). Conclusively, these results demonstrated that hydrogen generated from the 3S1C-Hyd-2 assembly effectively scavenged harmful ROS, reprogramed macrophages toward an anti-inflammatory phenotype, and enhanced cell survival under oxidative stress. This strategy established a promising platform for regulating inflammation via controlled hydrogen delivery.

### 2.5 Promotion of diabetic wound healing by the 3S1C-Hyd-2 assembly

Impaired diabetic wound healing arises from complex pathological interactions, with excessive ROS at lesion site playing a central regulatory role [42,43]. Targeted approaches that enable *in situ* ROS elimination and alleviate oxidative stress hold significant promise for chronic wound therapy. Given the potent anti-inflammatory and anti-apoptotic effects of 3S1C-Hyd-2 assemblies observed *in vitro*, we next explored their therapeutic efficacy *in vivo* by employing a streptozotocin (STZ)-induced diabetic mouse model (Fig. 5a, blood glucose levels exceeding 16.7 mmol L⁻¹ for seven consecutive days, as confirmed by the oral glucose tolerance test presented in Fig. S25). Full-thickness excisional wounds (7 mm) were created and treated with either Fmoc-KYF-AgNCs or Fmoc-KYF-AgNCs-3S1C-Hyd-2 hydrogels under light illumination. As shown in Fig. 5b, 3S1C-Hyd-2 assembly treatment markedly accelerated wound closure compared to controls, achieving ∼70% closure by day 7 and ∼87% by day 11, even surpassing the healing rates observed in non-diabetic mice. Histological evaluation at day 14 post-injury confirmed enhanced tissue regeneration in the Hyd-2 assembly group. Hematoxylin and eosin (H&E) staining revealed significantly thicker granulation tissue, while Masson’s trichrome staining showed denser collagen deposition and the reappearance of skin appendages such as hair follicles and sebaceous glands (Fig. 5c and S26).

**Fig. 5.**
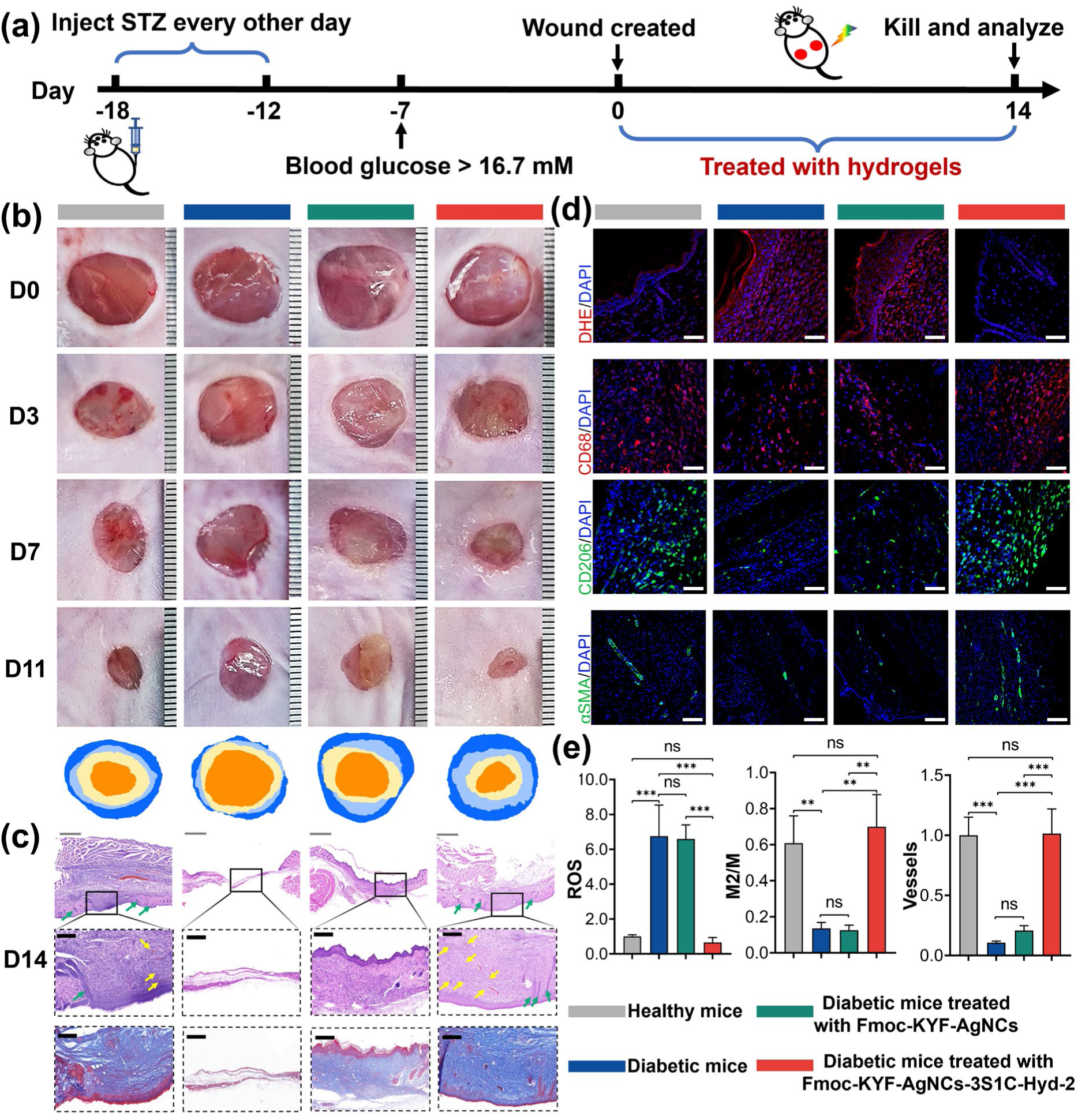
Therapeutic efficacy of Hyd-2 assembly in diabetic wound healing. (a) Schematic illustration of establishment of the mouse model for diabetic wound and the experimental design to evaluate the therapeutic efficacy of the Hyd-2 assembly. (b) Temporal progression of wound healing (Day 0/3/7/11) with representative macroscopic images. (c) Histopathological analysis of regenerated tissue of Day 14: Top – H&E staining showing granulation tissue and angiogenesis (scale bar: 1mm top; and 200 μm down.); Bottom – Masson trichrome staining quantifying collagen deposition (scale bar: 200 μm). (d) Multichannel immunofluorescence mapping: DHE/DAPI (ROS, red/nuclei), CD68/CD206/DAPI (M/M2 macrophages, red/green/nuclei), α-SMA/DAPI (neovascularization, green/nuclei), scale bar: 40 μm. (e) Quantitative comparisons of ROS intensity (left), M2/M polarization ratio (CD206^+^/CD68^+^, middle), and vascular density (α-SMA+ structures/mm², right). Data expressed as mean ± SD (n=5 biological replicates). Statistical significance: ns, no significance, *P<0.05, **P<0.01, ***P<0.001; one-way ANOVA with Tukey post-hoc.

To investigate the mechanistic basis of improved healing, we assessed ROS levels in wound tissues using the ROS-specific fluorescent dye dihydroethidium (DHE). Consistent with the recognized role of ROS in impairing diabetic wound repair, strong fluorescence signals were observed in untreated diabetic wounds and Fmoc-KYF-AgNCs-treated controls. In contrast, 3S1C-Hyd-2-assembly-treated wounds exhibited markedly reduced ROS accumulation (Fig. 5d, e), indicating effective ROS scavenging by the hydrogenase assemblies. As ROS levels critically regulate macrophage behavior, we further examined macrophage polarization by immunofluorescence staining for CD68 (pan-macrophage marker) and CD206 (M2 macrophage marker). Wounds treated with Hyd-2 assembly showed a substantial increase in CD206⁺ M2 macrophages compared to controls (Fig. 5d, e), suggesting enhanced resolution of inflammation and promotion of tissue remodeling. Furthermore, immunostaining for α-smooth muscle actin (α-SMA) demonstrated a significantly higher vessel density in the Hyd-2-assembly-treated group, indicative of improved angiogenesis critical for wound regeneration. Collectively, these findings demonstrated that the light-activated 3S1C-Hyd-2 assembly not only alleviated oxidative stress but also regulated macrophage polarization and enhance vascularization, thereby synergistically promoting diabetic wound healing and tissue regeneration.

To further clarify the potential mechanism of wound healing by the 3S1C-Hyd-2 assembly, we performed RNA sequencing (RNA-seq) of wound tissues from diabetic mice treated with 3S1C-Hyd-2 assemblies (Treated group) and untreated controls (diabetic group) at days 1, 7, and 14 post-treatments (Fig. S27-S29). Day 7 showed the most significant gene expression changes, with 2378 up-regulated and 2786 down-regulated genes (Fig. 6a). Given that ROS and inflammation are major barriers to healing, we performed Gene ontology (GO) and Kyoto Encyclopedia of Genes and Genomes (KEGG) enrichment analysis of down-regulated genes and revealed pathways associated with ROS production, immune responses, and inflammation were enriched (Fig. 6b, c and S30-S31). Notably, markers for M1 macrophages were down-regulated, whereas M2 markers were up-regulated (Fig. S32), which were consistent with our immunohistochemical results (Fig. 6d). Furthermore, we discovered that ROS metabolic processes, Toll-like receptor (TLR) signaling pathways, NF-κB signaling cascades, and NLRP3 inflammasome activity were significantly downregulated in both tissue and cell samples (Fig. S33), suggesting that Hyd-2 assembly can inhibit multiple inflammation-related pathways. To the up-regulated genes, functional enrichment analysis revealed that they were primarily involved in tissue regeneration, skin development, and wound healing (Fig. 6d and S34). Gene set enrichment analysis (GSEA) further confirmed the activation of pathways related to “skin epidermis development” and “keratinocyte differentiation” (Fig. S35), with significant up-regulation of keratin genes (*Krt1, Krt2, Krt5*) and other tissue repair genes (*Irf6, Cdh3, Krt9*) (Fig. 6e, f). Taken together, these findings suggest that 3S1C-Hyd-2 assemblies promote wound healing by clearing ROS, reducing inflammation, and enhancing tissue regeneration. The treatment facilitates M2 macrophage polarization, offering a promising approach for chronic wound management in diabetic conditions.

**Fig. 6.**
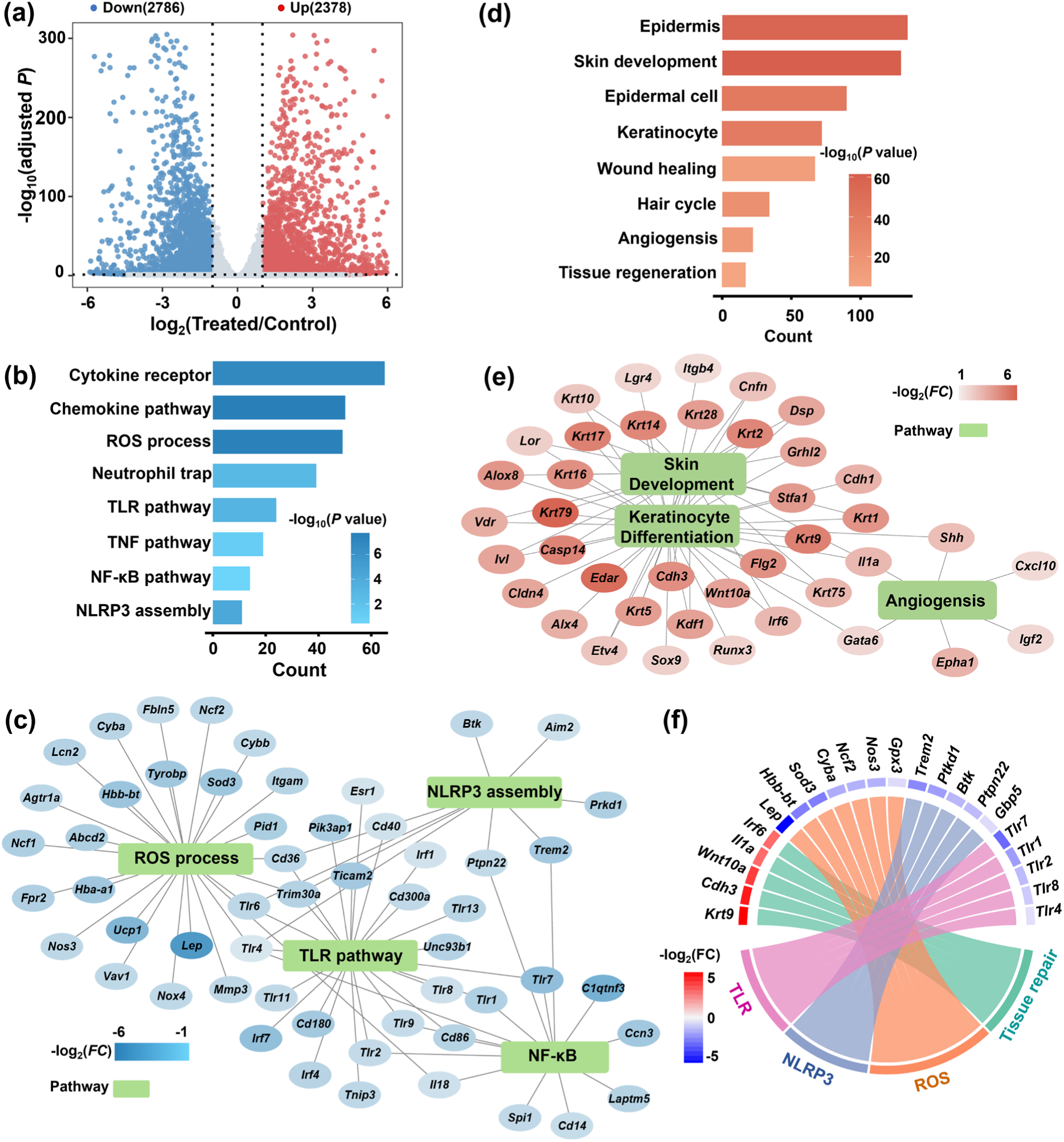
Transcriptomic analysis reveals biohydrogen-mediated modulation of inflammation and tissue repair in diabetic wounds on day 7. (a) Volcano plot depicting differentially expressed genes (DEGs) between the biohydrogen-treated and control groups. Red and blue dots represent significantly up- and down-regulated genes, respectively (adjusted *P* < 0.05). (b) Representative GO terms and KEGG pathways enriched in down-regulated genes, highlighting suppression of inflammation- and ROS-associated pathways. (c) Network of representative down-regulated DEGs enriched in inflammation-related GO terms; node color reflects log_2_(fold change). (d) Representative GO terms enriched in up-regulated genes showing activation of pathways related to tissue regeneration. (e) Network of representative up-regulated DEGs involved in wound healing processes; node color represents log_2_(fold change). (f) Chord diagram illustrating representative DEGs on day 7 associated with tissue repair, ROS metabolism, NLRP3 inflammasome, and Toll-like receptor signaling, indicating a coordinated transcriptional shift toward resolution of inflammation and promotion of repair.

Nature has evolved a variety of anaerobic enzymes—such as hydrogenases, nitrogenases [44], carbon monoxide dehydrogenases (CODH) [45], and formate dehydrogenases [46] —that exhibit exceptional catalytic properties, including high turnover frequency, remarkable substrate specificity, and negligible overpotential. However, these enzymes are inherently oxygen-sensitive, rendering them rapidly inactivated upon exposure to ambient oxygen, which severely limits their practical applications, particularly in biomedical contexts where aerobic conditions prevail. To harness their full catalytic potential in such settings, it is essential to establish engineered anaerobic microenvironments that effectively shield these enzymes from oxidative damage while maintaining their functional integrity. The design of the Fmoc-KYF-AgNCs hydrogel represents a paradigmatic synergistic strategy to address the critical challenge of oxygen sensitivity in hydrogenase applications, through a dual mechanism: physical barrier formation to restrict O_2_ diffusion and photoactivated AgNCs-mediated oxygen scavenging. The modular architecture of the peptide-AgNCs matrix enables integration with other oxygen-sensitive enzymes or therapeutic cargoes, offering a versatile platform for developing multifunctional therapeutics targeting complex disease systems.

Compared to existing systems, such as TiO_2_ nanorods for photocatalytic H_2_ generation [47] or algae–bacteria hydrogels requiring complex interspecies coordination [48], our system leverages the highly efficient hydrogen-evolving catalytic properties of hydrogenase. The [NiFe] active site selectively reduces H⁺ to H_2_ without generating harmful byproducts [49], operating effectively at physiological pH and temperature. This eliminates the cytotoxic risks associated with photothermal agents, such as hyperthermia-induced protein denaturation in AuPt@melanin hydrogels [50]. Notably, hydrogenases possess inherent catalytic reversibility (H_2_ ↔ 2H⁺ + 2e⁻), a feature rarely leveraged in therapeutic settings. In our platform, this reversibility enables adaptive H_2_ generation and consumption, supporting redox homeostasis under oxidative stress. Such self-regulation is absent in conventional chemical catalysts, including magnesium-based [51] or the tungsten bronze phase H_0.53_WO_3_-based [52] hydrogen release systems. This bidirectional activity better matches the dynamic physiological demands of wound healing—scavenging ROS during inflammation and supplying reducing equivalents during tissue remodeling. The 3S1C-Hyd-2 system applies precision medicine principles to address the spatiotemporal complexity of diabetic wounds. Unlike systemic H_2_ delivery (e.g., hydrogen-rich water), it confines H_2_ generation to the wound bed, minimizing off-target effects. Visible-light activation enables on-demand, adjustable H_2_ release, providing dynamic therapeutic control. Beyond ROS scavenging, H_2_ exerts immunomodulatory effects—suppressing proinflammatory genes (*Tnfα, Il1b*), promoting M2 macrophage polarization, and accelerating inflammation resolution, as confirmed by RNA-seq analysis. Upregulated α-SMA further indicates enhanced revascularization, counteracting ischemic hypoxia. Compared with conventional delivery, this localized enzymatic strategy maintains therapeutic H_2_ levels, offering superior bioavailability over gas inhalation or saline injection. This self-regulating catalytic mechanism provides a blueprint for extending reversible biocatalyst platforms (e.g., CODH) to broader gasotransmitter-based therapies.

## 3. Conclusion

In summary, we established a hybrid peptide–nanocluster hydrogel that creates a self-sustained anaerobic microenvironment, enabling the stable catalytic function of hydrogenase under aerobic or physiological conditions. Through the synergistic integration of physical confinement and photoactivated oxygen scavenging, this system effectively overcomes the intrinsic oxygen sensitivity of hydrogenases, achieving continuous light-driven hydrogen production in air. The generated hydrogen acts as a selective antioxidant to neutralize cytotoxic reactive oxygen and nitrogen species, thereby mitigating inflammation, promoting M2 macrophage polarization, enhancing angiogenesis, and accelerating tissue regeneration in diabetic wounds. Our work not only introduces a generalizable strategy for protecting oxygen-labile metalloenzymes but also pioneers the application of hydrogenase catalysis in biomedical therapy. The light-responsive, self-regulating mechanism allows spatiotemporal control of hydrogen generation, providing a dynamic approach to restore redox homeostasis and modulate immune responses. Beyond diabetic wound healing, this modular material–enzyme platform offers a versatile blueprint for extending anaerobic enzymatic catalysis toward other oxidative stress–related diseases and bioelectrocatalytic medical systems. Thereby, our study establishes a new paradigm for engineering functional anaerobic niches through synergistic material–enzyme–light interactions, bridging the gap between fundamental enzymology and therapeutic translation.

## Conflict of interest

The authors declare that they have no conflict of interest.

## Acknowledgment

Research reported in this publication was supported by the National Key Research and Development Program of China (No.2020YFA0907300).

## Author Contributions

1. L. Y. Z.: project administration, funding acquisition, methodology, investigation, formal analysis, data curation, conceptualization, writing—review and editing, and writing—original draft. C. Q. Z., B. F. S and Q.P.: methodology, investigation, formal analysis, data curation and writing—review and editing. H. S. J, W. J. L, Q. X, H. L, Y. X. G and Y. L. W: methodology, investigation, data curation and writing original draft. J. J. L. and Q.P.: theoretical calculation and molecular dynamics simulation. K. X. Y., C. X. X., X. Y. L. and B. F. S.: bioinformatics analysis and writing—review and editing. W. Z. L. C. J. and T. P. L.: visualization, discussion, writing—review and editing.

## Supporting Information

Any methods, additional references, Nature Portfolio reporting summaries, source data, extended data, supplementary information, acknowledgements, peer review information; details of author contributions and competing interests; and statements of data and code availability are available in supplementary materials.

## Notes

### Competing Interest Statement

The authors have declared no competing interest.

