## Supplemental Materials for "Engineering an Anaerobic Microenvironment to Empower Hydrogenase-Catalyzed Hydrogen Therapy for Diabetic Wound Healing"

* Corresponding authors.

### 1. Methods

#### 1.1 Materials

Polymethacrylic acid (PMAA, 674004), $AgNO_3$ (85228), tetrahydrofuran (THF, 401757) dimethyl formamide (DMF, 319937), methylene chloride (DCM, 270997), 2-(7-Azabenzotriazol-1-yl)-N,N,N',N'-tetramethyluronium hexafluorophosphate (HATU, 445460), N,N-diisopropylethylamine (DIPEA, D125806), lipopolysaccharides (LPS, L2630), dihydroethidium (DHE, D7008), 2',7'-dichlorofluorescein diacetate (H2DCFDA, 287810), and streptozotocin (STZ, 572201) were purchased from Sigma-Aldrich. Haematoxylin & eosin stain kit (G1120) and Masson's trichrome stain kit (G1340), and TriQuick total RNA extract reagent (R1100) were obtained from Solarbio. Hifair II 1st Strand cDNA Synthesis SuperMix (11123ES60) and Hieff UNICON qPCR SYBR Green Master Mix (11198ES08) were purchased from Yeasen. Antibodies CD68 (ab125212), CD206 (ab64693), α-SMA (ab7817), Alexa Fluor 488 (ab150113), Alexa Fluor 594 (ab150080) were purchased from Abcam.

#### 1.2 Enzyme preparation

The 3S1C-Hyd2 hydrogenases were produced as described previously [1,2]. In brief, the 3S1C-Hyd-2 protein was expressed in *Escherichia coli* K-12 strain HJ001-hyp-3S harboring the corresponding plasmid and subsequently purified through a multi-step protocol. This included Ni-affinity chromatography, desalting via PD-10 column chromatography, and final purification by size exclusion chromatography on a Superdex 200 column. The collected protein fractions were assessed for purity using SDS-PAGE. Protein concentration was quantified with the Bradford assay kit, and the purified Hyd-2 protein was stored under liquid nitrogen conditions for future applications.

#### 1.3 Synthesis of silver nanoclusters (AgNCs)

AgNCs were prepared as described previously [3,4]. In the general protocol, $Ag^+$ ions and PMAA were introduced into an aqueous solution at a molar ratio of 2:1 ($Ag^+$: MAA). The solution was subsequently degassed by purging with $N_2$ for 30 minutes. Exposure to solar radiation for 9 hours facilitated the formation of AgNCs. The resulting AgNCs were purified via precipitation using a 17-30% (v/v) THF solution under vigorous stirring. The precipitate obtained after centrifugation was confirmed to consist of AgNCs.

#### 1.4 Synthesis of Fmoc-KYF peptide

The tripeptide Fmoc-KYF (Fmoc-Lys-Tyr-Phe) was synthesized using standard Fmoc-based solid-phase peptide synthesis (SPPS) on CLEAR amide resin (loading: 0.1 mmol) [5,6]. The resin was swollen in DMF for 30 min and sequentially washed with DMF, DCM, and DMF. Fmoc deprotection was performed using 20% (v/v) piperidine in DMF, with a brief 1 min pre-rinse followed by a 20 min incubation, then washed as above. Each coupling cycle employed 8 equivalents of the respective Fmoc-protected amino acid, preactivated in 0.5 M HATU and DIPEA (0.8 mmol each) in DMF. Coupling proceeded for 25 min per residue. After chain assembly, the peptide remained N-terminally Fmoc-protected.

Cleavage from the resin was achieved by treatment with TFA/$H_2O$/TIS (95:2.5:2.5, v/v) for 3 h at room temperature. The cleavage solution was collected, and residual peptide was washed from the resin with additional TFA mixture. The combined eluates were concentrated under reduced pressure and precipitated with cold diethyl ether (−20 °C). The precipitate was collected by centrifugation (4000 g, 20 min), washed three times with ether, and air-dried overnight. The crude product was dissolved in 30% acetonitrile/$H_2O$ (v/v) and lyophilized. Purity was confirmed by analytical RP-HPLC using a C18 column, and the molecular mass was verified via MALDI-TOF mass spectrometry. The structure was confirmed by $^1$H NMR and $^{13}$C NMR spectroscopy, using an AVANCE NEO 400 MHz spectrometer (Bruker).

To prepare a homogeneous hydrogel, Fmoc-KYF was dissolved in water at a concentration of 10 mg mL$^{-1}$ to form a stock solution. AgNCs or $Na^+$ ions were employed to promote gelation. For hydrogel formation in vials, 500 μL of the Fmoc-KYF stock solution was transferred into a vial, followed by the addition of 500 μL of AgNCs solution (120 μM) or 200 mM $NaNO_3$ solution. The mixture was then homogenized by thorough mixing and incubated for several minutes to allow gelation. For electrode-based applications, 5 μL of the Fmoc-KYF stock solution (10 mg mL$^{-1}$ in water) was deposited onto a PGE electrode. Subsequently, 5 μL of AgNCs solution (120 μM) or 5 μL of 200 mM $NaNO_3$ solution was added. After gentle mixing, the mixture was incubated for several minutes to facilitate gelation.

### 1.5 Transmission electron microscopy Imaging (TEM)

The Fmoc-KYF assemblies were prepared at a concentration of 100 μM and adsorbed onto carbon-coated, 300-mesh copper grids (Zhongjingkeyi Technology Co. Ltd., Beijing, China; glow-discharged before use) for 2 minutes. Excess solution was removed using the filter paper. After drying, TEM micrographs were recorded on a Talos F200X G2 transmission electron microscope at an acceleration voltage of 120 kV.

### 1.6 Atomic force microscope imaging (AFM)

Samples of Fmoc-KYF hydrogel or Fmoc-KYF-AgNCs hydrogel were prepared and incubated on freshly cleaned mica substrates for 10 minutes. Subsequently, the samples were thoroughly rinsed multiple times with deionized water to remove any non-adherent material. Imaging was performed in liquid phase using tapping mode. A probe model SNL-10-B with a resonance frequency of approximately 5 Hz was utilized for this purpose. All atomic force microscopy (AFM) measurements were conducted using a BRUKER MultiMode 8 AFM system, and representative micrographs were recorded.

### 1.7 Circular dichroism spectroscopy

CD spectra of Fmoc-KYF were acquired at a concentration of 5 mg $mL^{-1}$ using a J-810 CD spectropolarimeter (JASCO, Japan) equipped with a 0.1 cm quartz cell. Spectra were recorded in the wavelength range of 190 to 260 nm under the following conditions: a scan speed of 200 nm $min^{-1}$, a step resolution of 2 nm, and three accumulations at room temperature.

### 1.8 Fluorescence spectroscopy

For fluorescence spectroscopy experiments, Fmoc-KYF was prepared as a 14 mM aqueous solution, whereas AgNCs were dissolved in ultrapure water to achieve a concentration of 1 mM and used as a stock solution. Both solutions were subsequently diluted with deionized water for further spectroscopic analyses. Fluorescence spectra were acquired using a fluorescence spectrophotometer (model FPro970). The emission spectra were recorded at an excitation wavelength of 460 nm for AgNCs and 280 nm for the Fmoc-KYF peptide.

### 1.9 Absorbance spectroscopy

For absorbance spectroscopy experiments, Fmoc-KYF was prepared as a 14 mM aqueous solution, whereas AgNCs were dissolved in ultrapure water at a concentration of 1 mM to serve as a stock solution. Both solutions were subsequently diluted to various concentrations using water for further testing with a UV-vis spectrophotometer (U-3900, Hitachi High-Tech Group).

### 1.10 FTIR spectroscopy

The hydrogels were prepared as the described in the gelation part, then freeze-dried. 4 mg of freeze-dried material mixed with 2 g of potassium bromide was ground into powder, pressed into thin sheets and tested on the Fourier transform infrared spectrometer (Tensor 37, Bruker).

### 1.11 DFT calculations

High precision molecular conformation search was performed using Molclus (Version 1.12, http://www.keinsci.com/research/molclus.html) combined with xtb program [7] to generate molecular dynamics trajectories using GFN0-xTB method [8], then perform batch optimization under the GFN2-xTB method [9]. DFT calculations were performed for a batch of low-energy conformations obtained through the above conformational search. All the conformers discussed in the work were optimized using the Gaussian 09 suite of programs [10] associated with the B3LYP-D3 [11] /6-31G(d, p) level [12,13]. Solvation effects of water were taken into account by performing optimization using the SMD model [13]. Stationary points were confirmed with vibrational frequency computations, with ground states having zero imaginary frequency.

### 1.12 MD simulations

The OPLS-AA atomic force field [14] was chosen for the simulation, and the Gromacs2018.8 software package [15] was used for the molecular dynamics (MD) calculations. The potential functions include bonding potentials such as bond lengths, bond angles and dihedral angles, and non-bonding interaction potentials, where the non-bonding interactions include the Lennard-Jones potential (for van der Waals interactions) and the Coulomb potential (for electrostatic interactions).

The initial structure of Fmoc-KYF hydrogels corresponds to the DFT calculated optimal structure (Fig. S15) of the fibrils containing 44 assembled Fmoc-KYF peptides. The dimensions of the initial structure of the Fmoc-KYF fibrils were 50×45×20 Å. In the respective simulation systems (Fig. S16), twenty $O_2$, or $H_2$ molecules were initially positioned in a cubic 70×70×70 Å water box surrounding the Fmoc-KYF fibril.

After the initial configurations were established, energy minimization procedures and a 1 ns equilibration phase were carried out prior to performing ten replicate 50 ns molecular dynamics (MD) simulations under the NPT ensemble. The simulation step size was 2 fs, and the kinetic trajectory information was saved every 100 ps for subsequent analysis, and the VMD program was used to observe the kinetic trajectories. The water molecules were modeled by simple point charge (SPC) model [16], and the temperature control method was adopted by Velocity-rescale thermal bath method [17]. Periodic boundary conditions were used in the three-dimensional direction, and the LINCS algorithm was used to constrain the bond lengths of the molecules, and the Particle-mesh Ewald (PME) method was chosen to deal with the long-range electrostatic interactions, and the truncation radius of non-bonded interactions was set to 1.0 nm.

#### 1.13 Oxygen scavenging test

For gas control, three tubes of Fmoc-KYF (10 mg $mL^{-1}$) and three tubes of AgNCs solutions (120 μM containing 20 μM $H_2DCFDA$) were sparged with pure $H_2$, $O_2$, or $N_2$, respectively. Subsequently, the solutions were mixed in varying volumes to form hydrogels with different gas concentrations. The gelation process was conducted as described in the gelation section, resulting in a final concentration of 5 mg $mL^{-1}$ for Fmoc-KYF, 60 μM for AgNCs, and 10 μM for $H_2DCFDA$. For oxygen scavenging, Fmoc-KYF-AgNCs hydrogel was prepared under an atmosphere of 21% $O_2$ and 79% $N_2$. Following gelation, the sample was exposed to illumination for 120 seconds using a 300 W Arc Lamp (Newport 69911), equipped with a 420 nm filter and positioned 10 cm from the cuvette. The fluorescence of the ROS-specific probe $H_2DCFDA$ was measured using a fluorescence spectrophotometer (FPro970) at an excitation wavelength of 488 nm. For ROS scavenging, Fmoc-KYF and AgNCs solutions were sparged with a gas mixture of 21% $O_2$ and 79% $H_2$. All subsequent procedures were identical to those used in the oxygen scavenging experiments.

### 1.14 Protein film electrochemistry

All PFE experiments were conducted in an anaerobic glovebox (Mikrouna) under a nitrogen atmosphere with oxygen levels maintained below 3 ppm. Electrochemical measurements were performed using an Autolab potentiostat (model J101), controlled by Nova software (version 2.1). A three-electrode configuration was utilized, comprising a platinum wire as the counter electrode and an Ag/AgCl (3M) electrode as the reference electrode. The protein film was fabricated by depositing 3 μL of hydrogenase solution (10 μM) onto the working electrode via drop-casting. The buffer solution employed in all experiments consisted of 15 mM sodium acetate, 15 mM CHES, 15 mM MES, 15 mM HEPES, 15 mM TAPS, and 0.1 M $NaNO_3$ at pH 6.0. All potentials were converted to the Standard Hydrogen Electrode (SHE) scale using the correction $E_{SHE} = E_{Ag/AgCl} + 197$ mV at 25 °C.

### 1.15 Photocatalytic experiments and H2 quantification

For photocatalytic experiments, all samples were prepared inside a glovebox under a $N_2$ atmosphere before being removed for measurements. Briefly, the 3S1C-Hyd-2 hydrogenase was mixed with AgNCs in a molar ratio of 1:1 and incubated in the dark for 30 minutes to form a complex. This complex was then added evenly to 0.5 mL of a 10 mg $mL^{-1}$ Fmoc-KYF peptide aqueous solution, followed by the addition of 0.5 mL of AgNCs solution (120 μM) containing 0.1 M TEOA at pH 7.0. After gelation, 1 mL of triethanolamine (TEOA) buffer (0.1 M, pH 7.0) was added to the vial. The mixture was degassed and sealed, and removed from the glove box for photochemical experiments. For the aerobic group, air was purged into the vial prior to conducting photochemical experiments. For photocatalysis measurements, the samples were irradiated with visible light using a 300 W Arc Lamp, (Newport 69911) fitted with a 420 nm filter and held 10 cm from the vessel. Temperature was controlled by immersing the vessel in a water bath. Production of $H_2$ was monitored at regular intervals by removing small volumes (50 μL) of headspace gas for gas chromatography (GC) analysis. A Shimadzu GC-2030 series gas chromatograph (GC) with electronic pneumatic control, optimized for trace gas analysis, was used for $H_2$ quantification. Headspace gas samples of the reaction mixture were injected into the GC operating in pulsed splitless mode using a gas-tight Hamilton syringe. The chromatographic separation was performed using a Micropacked ST P/N MP-01 column (Serial No. 0341) with a barrier discharge ionization detector and high-purity He carrier gas were used.

#### 1.16 Cell experiments

The mouse macrophage cell line (RAW264.7) was obtained from the American Type Culture Collection (ATCC). The cells were maintained in Dulbecco's Modified Eagle Medium (DMEM) supplemented with 10% fetal bovine serum (FBS) under a humidified atmosphere of 5% $CO_2$ at 37°C. For hydrogel treatment, the 3S1C-Hyd-2 hydrogenase was combined with AgNCs at a molar ratio of 1:1 and incubated in the dark for 30 minutes to form a complex. This complex was subsequently mixed uniformly into 0.25 mL of a 10 mg $mL^{-1}$ Fmoc-KYF peptide aqueous solution within one well of a six-well plate. Following this, 0.25 mL of AgNCs solution (120 μM), containing 0.01 M TEOA at pH 7.0, was added. After gelation, a coverslip was gently placed on the surface of the hydrogel, followed by the addition of cell culture medium. Subsequently, $1 \times 10^6$ cells per well were seeded. Once the cells had adhered, the medium was replaced with fresh medium containing 100 ng $mL^{-1}$ lipopolysaccharide (LPS), and the cultures were incubated for 24 hours. During this period, the hydrogel located at the bottom of the six-well plate was irradiated with an LED lamp (14W, Bull). Finally, the cells were harvested for fluorescence staining or RNA extraction.

#### 1.17 CCK-8 test

Hydrogels were assembled as the cell experiments section in 96-well plates, and $5 \times 10^3$ cells per well were seeded with medium contains 100 ng $mL^{-1}$ LPS. After the cells were irradiated with an LED lamp (14W, Bull) in the cell culture incubator, the medium was replaced with new medium contains 10% CCK-8 detection solution. Place the 96-well plate in a cell culture incubator and incubate for 1 hour protected from light, then measure the absorbance at 450 nm with a microplate reader.

### 1.18 Real-time quantitative PCR (RT–qPCR)

Total RNA was extracted from RAW264.7 cells using the TriQuick Total RNA Extraction Reagent according to the manufacturer's protocol. The RNA concentration and purity were determined using a NanoDrop spectrophotometer (ND-100, MIULAB). Subsequently, 1 μg of total RNA was reverse-transcribed into cDNA using the Hifair II 1st Strand cDNA Synthesis SuperMix for qPCR. Real-time quantitative PCR (RT-qPCR) was performed with the Hieff UNICON qPCR SYBR Green Master Mix on a Bio-Rad CFX384 Real-Time PCR Detection System (Bio-Rad).

### 1.19 Study in diabetic mouse model

Male BALB/c mice were utilized to establish a streptozotocin (STZ)-induced diabetes model. Specifically, the mice received intraperitoneal injections of STZ at a dose of 45 mg $kg^{-1}$ for five consecutive days. Two weeks post-injection, mice with blood glucose levels consistently ≥16.7 mmol $L^{-1}$ across two consecutive measurements were classified as diabetic [18]. Diabetic mice were subsequently anesthetized using 1% sodium pentobarbital, and two full-thickness skin wounds (7 mm in diameter) were created on their backs using a circular punch biopsy tool. Following wound creation, the mice were treated with one of the following: normal saline, Fmoc-KYF-AgNCs hydrogel, or Fmoc-KYF-AgNCs-3S1C-Hyd-2 hydrogel. The wounds were then covered with a transparent medical dressing and exposed to white light irradiation (30W, GLTX) for a duration of 2 hours. Images of the wounds were captured at various time points, and the wound areas were quantified using ImageJ software. The relative wound area was calculated according to the formula: relative wound area = wound area on a given day / wound area on day 0. RNA was extracted from tissue samples collected within 1 mm of the wound edge.

Gelation Details: All stock solutions of Fmoc-KYF (10 mg $mL^{-1}$) and AgNCs (120 μM containing 0.01 M TEOA, pH 7.0) were degassed for 30 minutes using $N_2$ prior to their application on wounds. For the preparation of the Fmoc-KYF-AgNCs hydrogel, 50 μL of Fmoc-KYF was applied to the wound, followed by the addition of 50 μL of AgNCs solution to induce gelation. In the case of the Fmoc-KYF-AgNCs-3S1C-Hyd-2 hydrogel, the 3S1C-Hyd-2 was mixed with AgNCs at a molar ratio of 1:1 and incubated in the dark for 30 minutes to form a complex. This complex was subsequently homogeneously incorporated into 50 μL of Fmoc-KYF peptide solution on the diabetic wound, after which 50 μL of AgNCs solution was added to facilitate gelation.

**1.20 Histology and immunofluorescence analysis**

Skin tissues were fixed in 4% paraformaldehyde, embedded in paraffin, and sectioned into 6-μm-thick slices for histological staining. Re-epithelialization was assessed using hematoxylin and eosin staining, while collagen deposition was evaluated with Masson's trichrome staining. ROS levels in wound tissues were quantified using the dihydroethidium (DHE) staining method. For immunofluorescence analysis, paraffin-embedded sections were deparaffinized in xylene and rehydrated through a graded ethanol series, followed by permeabilization with 0.1% Triton X-100 for 15 minutes. The samples were then incubated with primary antibodies overnight at 4°C. After six washes with phosphate-buffered saline (PBS), the samples were incubated with secondary antibodies for 2 hours at room temperature. Subsequently, the samples were counterstained with 4',6-diamidino-2-phenylindole (DAPI) after additional PBS washes. Finally, images were acquired using a total internal reflection fluorescence microscope (Leica DMi8).

**1.21 RNA-sequencing data analysis**

Library construction using Hieff NGS™ MaxUp Dual-mode mRNA Library Prep Kit for Illumina® (12301ES96, Yeasen) and sequencing were performed using DNBSEQ-T7 by Sangon Biotech. Quality control of raw sequence data was performed with trim-galore (v0.6.10) and checked using FastQC (v0.11.9), followed by alignment to the mouse genome mm10 using hisat2 (v2.2.1) [19]. Samtools (v1.9) was used to filter and sort the mapped reads [20], then featureCounts (v2.0.6) was used to summarize the gene counts [21]. Differential gene expression analysis utilized the DESeq2 package (v1.40.2) [22] with a cut off $\log_2$(fold change) > 1 and adjusted $P$ value < 0.05. Gene Ontology (GO) and Kyoto Encyclopedia of Genes and Genomes (KEGG) enrichment analysis of differentially expressed genes was performed using the cluster Profiler R package (v4.8.3) [23], and visualized by Cytoscape (v3.10.2) [24]. Gene Set Enrichment Analysis (GSEA) was applied to identify predefined gene sets displaying statistically significant differences between two given groups (http://www.broadinstitute.org/gsea/index.jsp) [25].

#### 1.22 Data availability

All generated or analyzed data supporting the findings of this study are available within the paper and its Supplementary Information. The RNA-seq data are available from the Gene Expression Omnibus (GEO) database, with accession number GSE297828. All raw data from this study are available from the corresponding authors upon request. Source data are provided with this paper.

#### 1.23 Statistics

All results are expressed as mean ± s.e.m. unless specified otherwise. Statistical analyses were performed using GraphPad Prism 8 (GraphPad Software). * denotes $P < 0.05$, ** denotes $P < 0.01$, *** denotes $P < 0.001$ and **** denotes $P < 0.0001$.

#### 1.24 Ethics oversight

All animal care and experimental procedures adhered strictly to the guidelines set forth in the National Institutes of Health (NIH) Guide for the Care and Use of Laboratory Animals. The mouse experiments were formally approved by the Institutional Animal Care and Use Committee (IACUC) of Nankai University, under the approved protocol number IACUC #2023-SYDWLL-000640.

### 2. SI Figure and tables

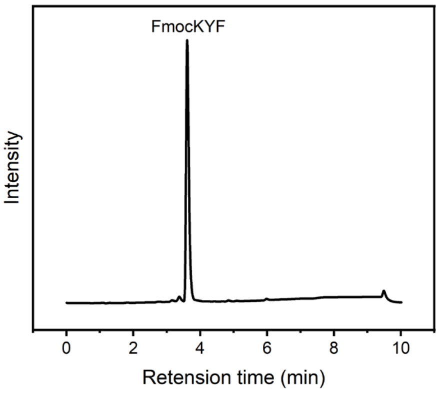


**Fig. S1**. High-Performance Liquid Chromatogram (HPLC) of the Fmoc-KYF peptide. Using a C18 column, the mobile phase A consisted of 100% methanol with 0.1% TFA, while mobile phase B comprised 95% water, 5% methanol, and 0.1% TFA. Detector 1 was set at 220 nm, and detector 2 at 254 nm.

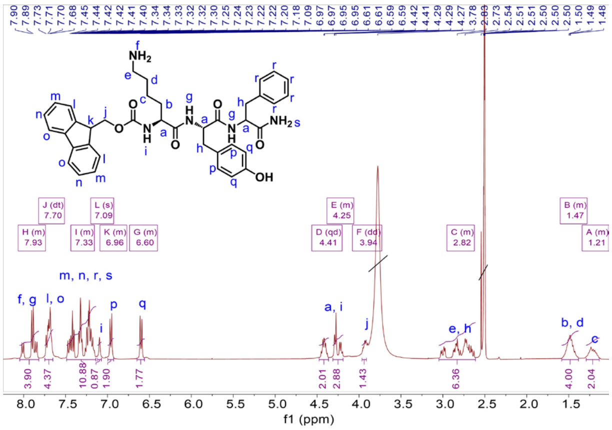


**Fig. S2.** NMR $^{1}$H spectrum of the peptide Fmoc-KYF. 400 MHz, DMSO-d6: δ 8.05 – 7.82 (m, 4H), 7.70 (dt, J = 14.1, 7.1 Hz, 4H), 7.47 – 7.40 (m, 2H), 7.32 (t, J = 1.7 Hz, 2H), 7.27 – 7.16 (m, 5H), 7.09 (s, 1H), 7.00 – 6.93 (m, 2H), 6.65 – 6.56 (m, 2H), 4.41 (qd, J = 7.9, 4.9 Hz, 2H), 4.32 – 4.18 (m, 3H), 3.94 (dd, J = 8.8, 5.1 Hz, 2H), 3.05 – 2.61 (m, 6H), 1.58 – 1.37 (m, 4H), 1.29 – 1.13 (m, 2H)。

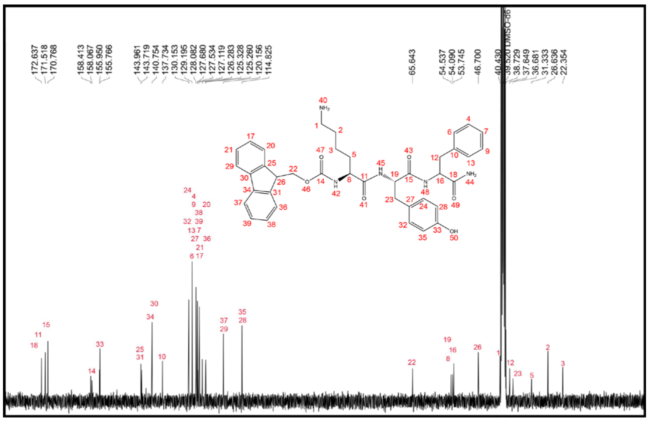


**Fig. S3.** NMR $^{13}$C spectrum of the peptide Fmoc-KYF. 101 MHz, DMSO-$d_6$: δ = 172.6, 171.5, 170.8, 158.4, 158.1, 155.9, 155.8, 144.0, 143.7, 140.8, 137.7, 130.2, 129.2, 128.1, 127.7, 127.5, 127.1, 126.3, 125.3, 125.3, 120.2, 114.8, 65.6, 54.5, 54.1, 53.7, 46.7, 40.4, 38.7, 37.6, 36.7, 31.3, 26.6, 22.4 ppm.

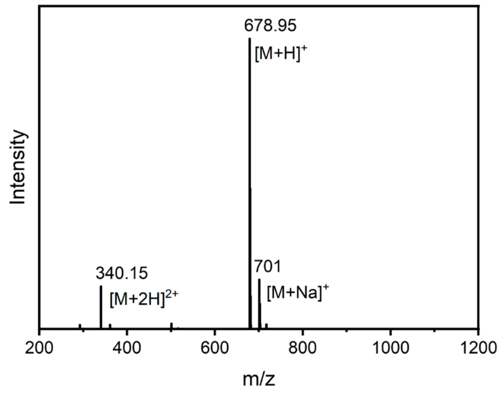


**Fig. S4.** Mass spectrum of the peptide Fmoc-KYF. M means Fmoc-KYF.

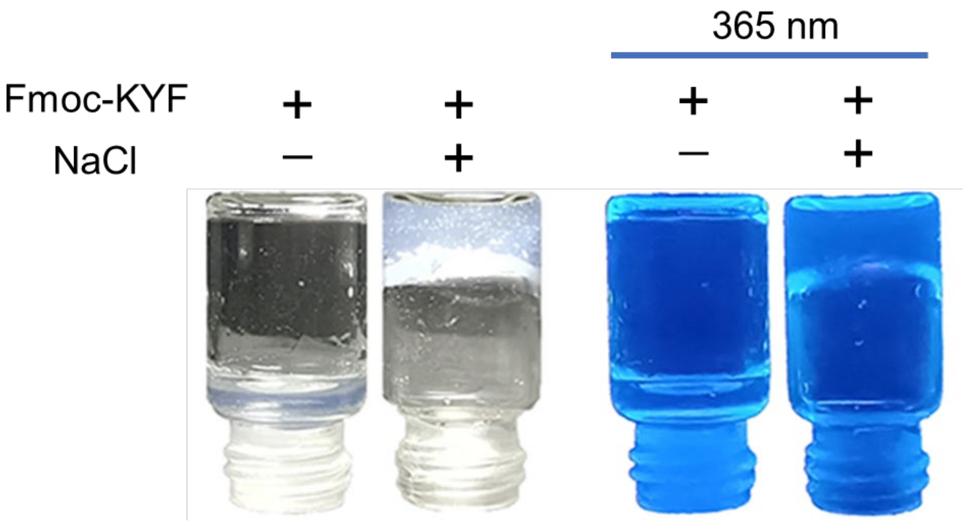


**Fig. S5**. The images of Fmoc-KYF hydrogel formation induced by NaCl.

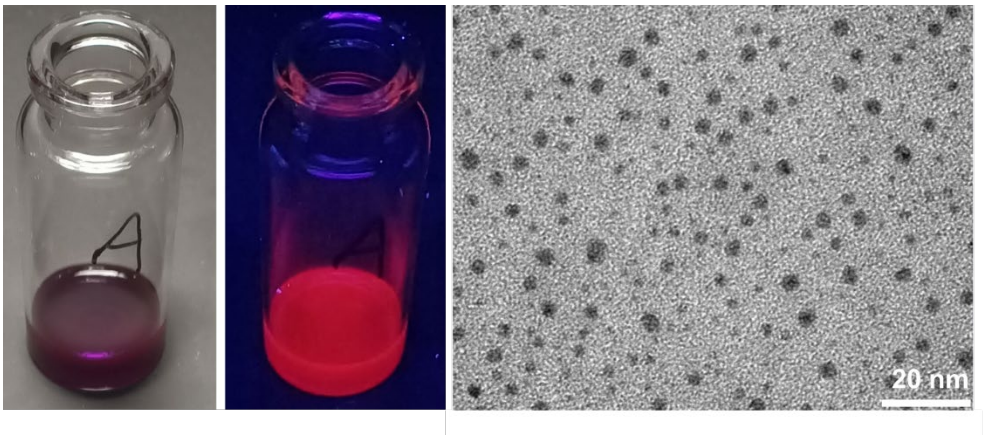


**Fig. S6**. The left image shows the photograph of the as-prepared AgNCs. The middle image displays the photograph of the as-prepared AgNCs under 365 nm irradiation. The right image presents the TEM image of the as-prepared AgNCs.

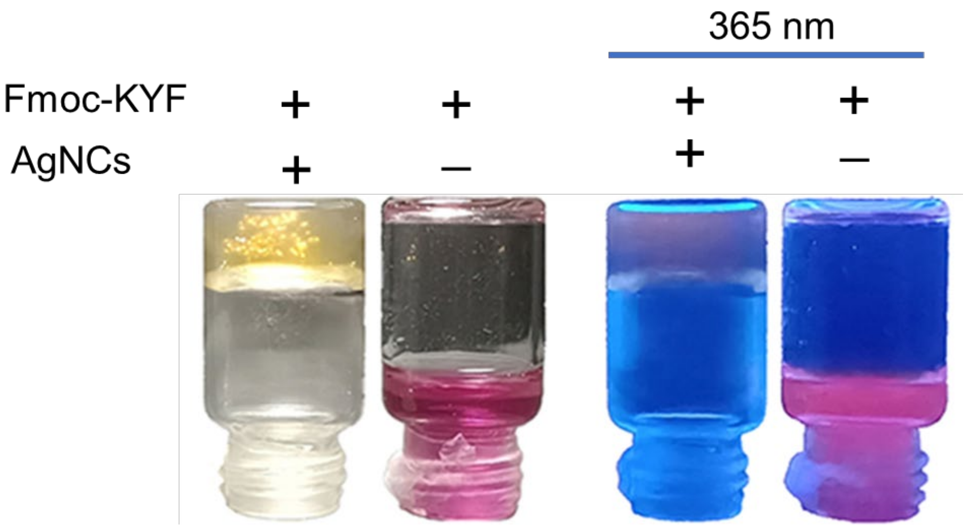


**Fig. S7**. The images of Fmoc-KYF-AgNCs hybrid hydrogel.

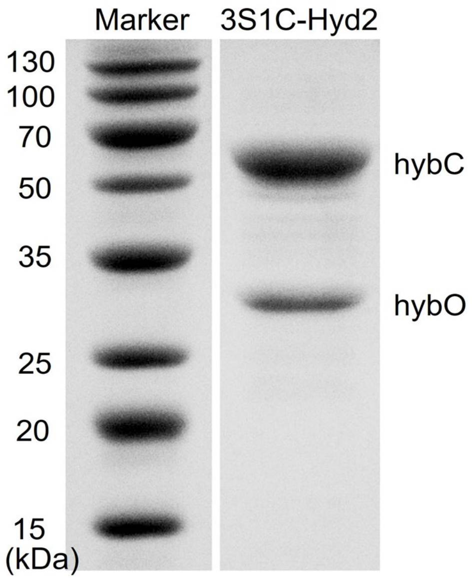


**Fig. S8.** SDS-PAGE analysis of purified 3S1C-Hyd-2 enzyme.

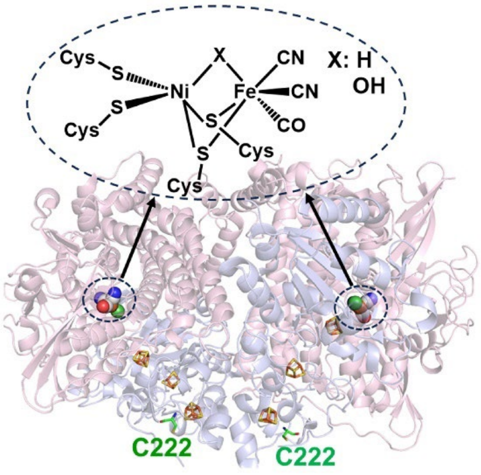


**Fig. S9**. Structure of 3S1C-Hyd-2 (PDBID: 6G7M) with bimetallic active site enlarged. The structure of the surface-exposed cysteine 222 residue is highlighted in green.

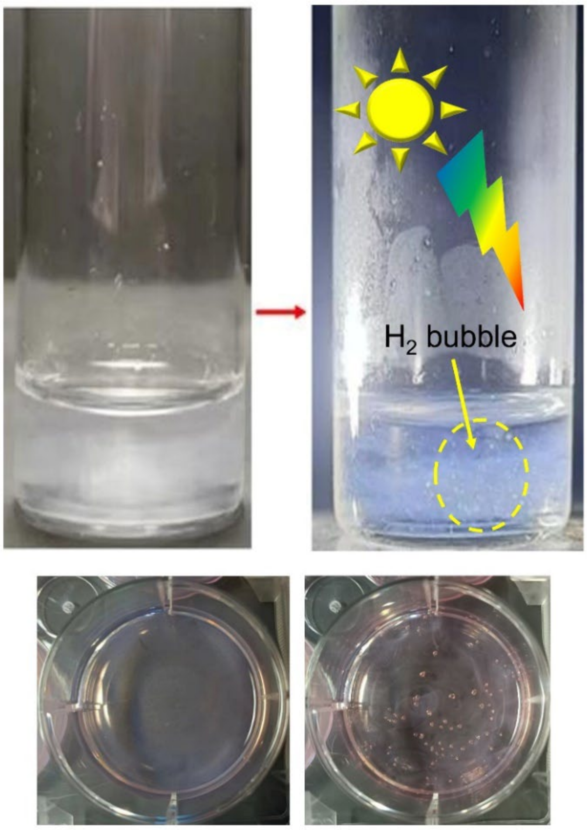


**Fig. S10**. The images illustrate the photocatalytic $H_2$ evolution performance of the Fmoc-KYF-AgNCs-3S1C-Hyd-2 system, whether in a glass vial (top) or within the wells of a cell culture plate(bottom).

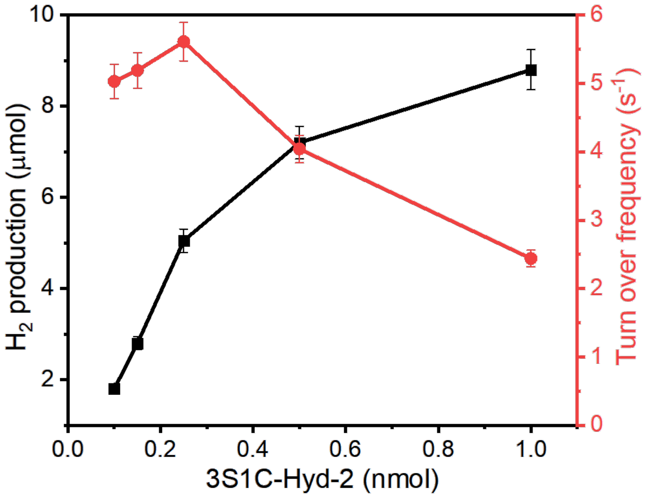


**Fig. S11.** Optimization of enzyme concentration for enhanced $H_2$ production. 2 mL total volume containing 0.1-1.0 nmol 3S1C-Hyd-2, 5 mg $mL^{-1}$ Fmoc-KYF, 0.5 mg $mL^{-1}$ AgNCs, 0.1 M TEOA, pH 7.0 under irradiation 2 hours.

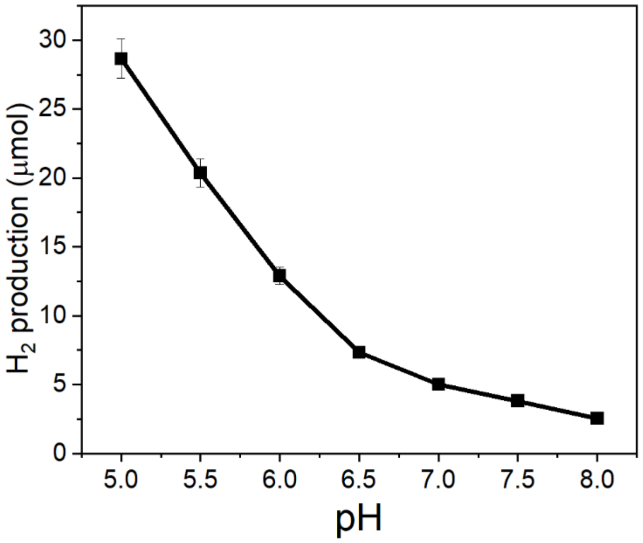


**Fig. S12.** The effect of pH on $H_2$ production. A total reaction volume of 2 mL containing 0.25 nmol of 3S1C-Hyd-2, 5 mg mL$^{-1}$ Fmoc-KYF, 0.5 mg mL$^{-1}$ AgNCs, and 0.1 M TEOA was prepared, with the pH adjusted between 5.0 and 8.0.

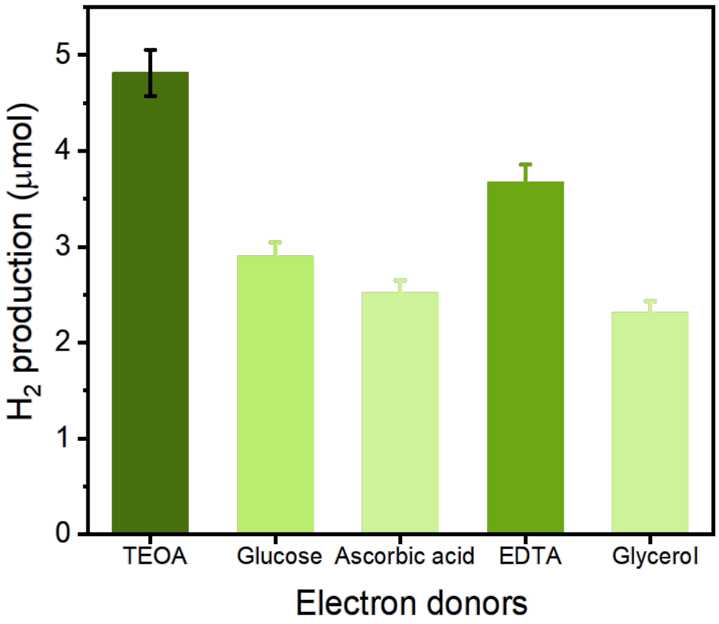


**Fig. S13.** The effect of electronic sacrificial agent on $H_2$ production. A total reaction volume of 2 mL containing 0.25 nmol of 3S1C-Hyd-2, 5 mg mL$^{-1}$ Fmoc KYF, 0.5 mg mL$^{-1}$ AgNCs, and 0.1 M electronic sacrificial agent (TEOA, glucose, Ascorbic acid, EDTA and glycerol) was prepared, pH 7.0.

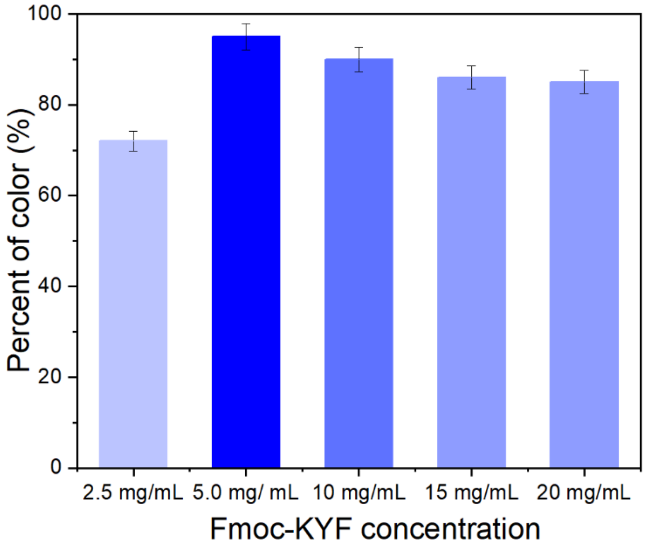


**Fig. S14.** Effect of Fmoc-KYF gel concentration (2.5- 20 mg $mL^{-1}$) on the fading of methyl viologen ($MV^{+\bullet}$), following a 3 hours exposure to atmospheric $O_2$ (n = 3).

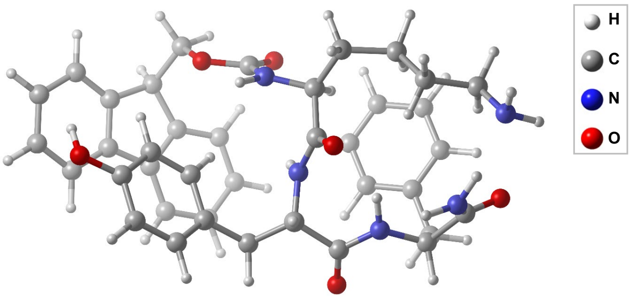


**Fig. S15.** The optimal structure of Fmoc-KYF.

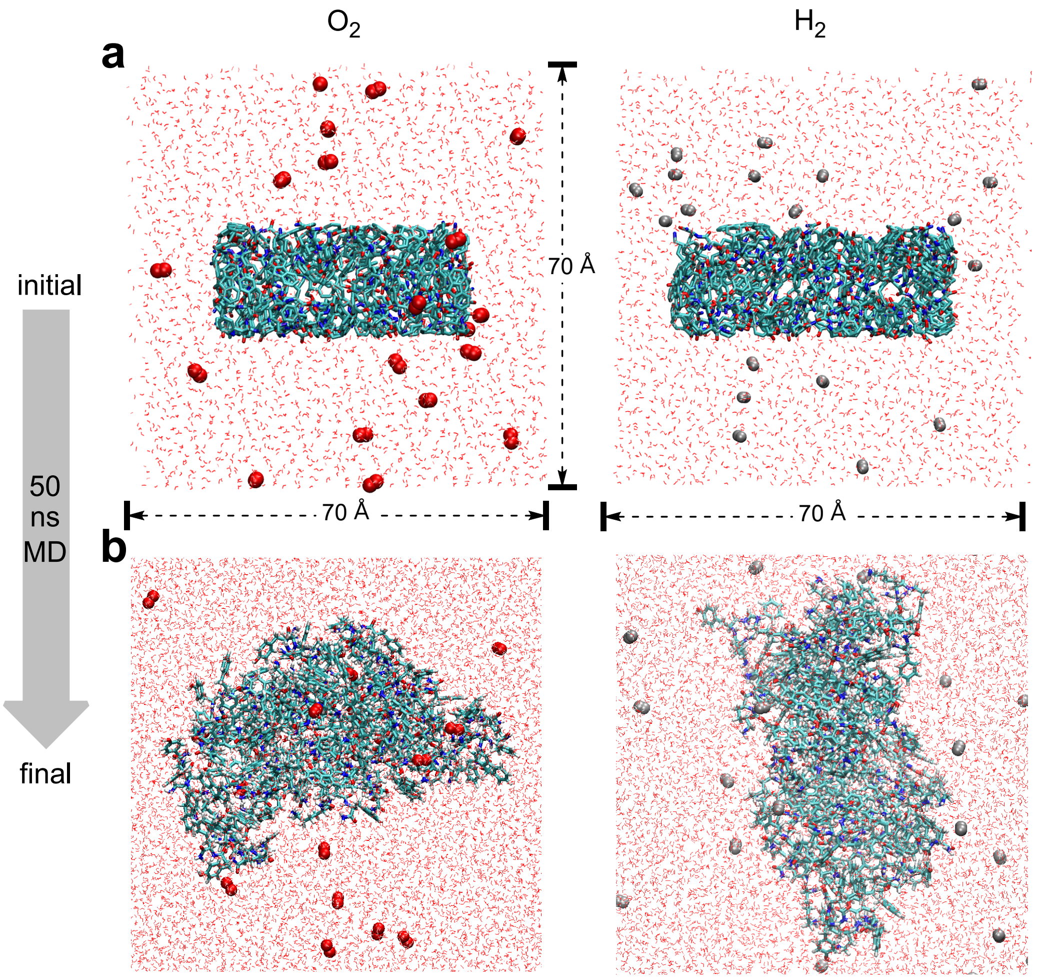


**Fig. S16.** Molecular graphics representations of the simulation system. (a) The initial simulation system. (b) The final simulation system after 50 ns MD. The simulated Fmoc-KYF fibril segment embedded in a 70 Å³ water box, showing $O_2$ (red color) and $H_2$ (silver color) molecules represented in van der Waals (VDW) format.

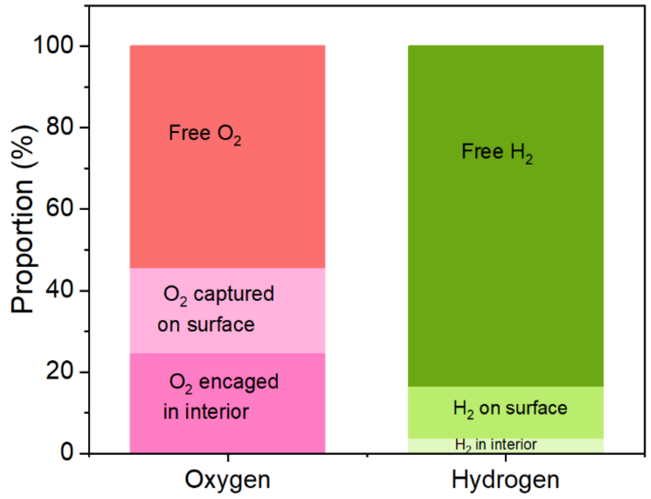


**Fig. S17**. Percentage of free and bound oxygen and hydrogen molecules in the presence of Fmoc-KYF fibrils.

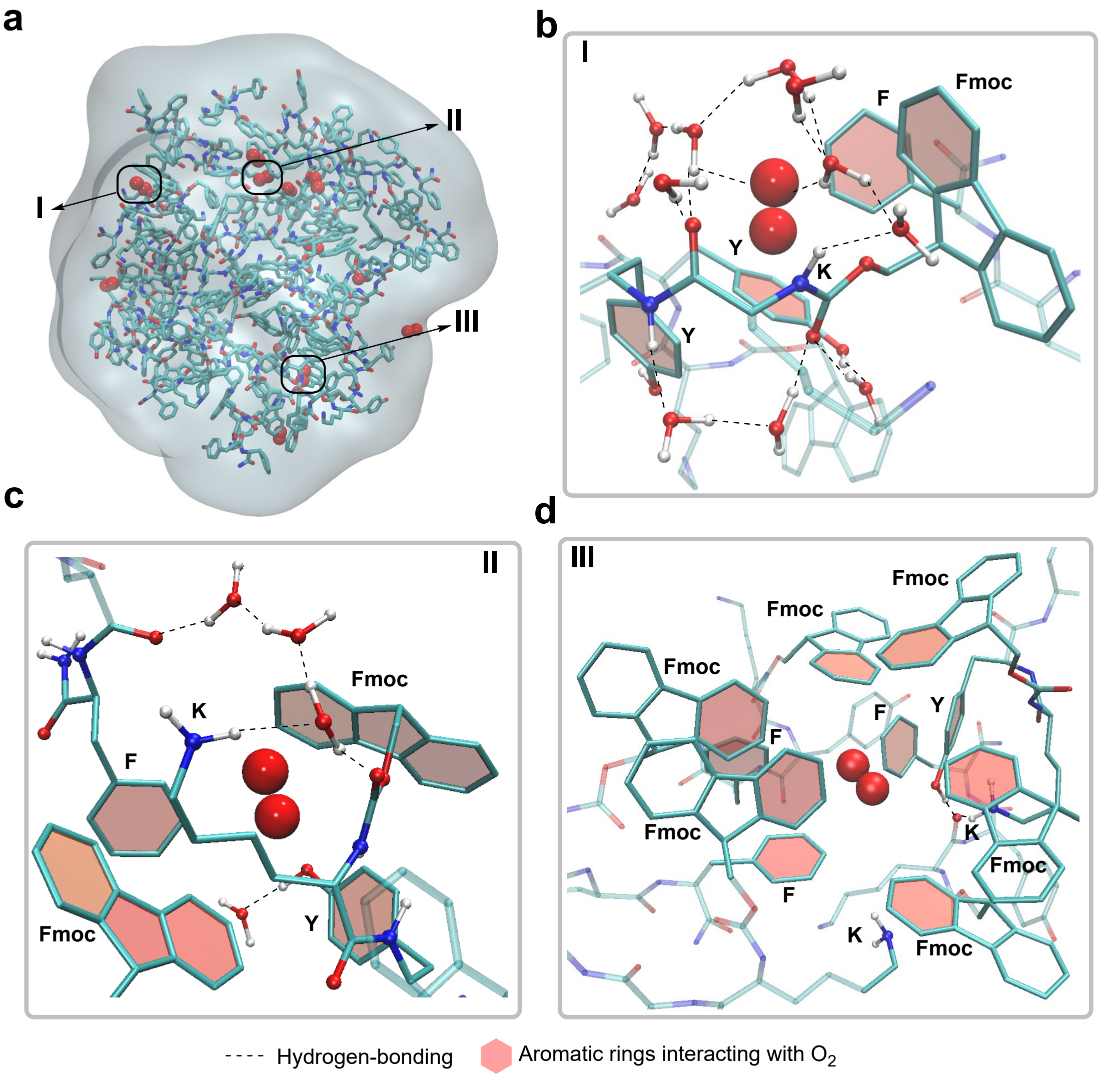


**Fig. S18.** Molecular dynamics simulation of $O_2$ binding to Fmoc-KYF hydrogels. (a) Molecular graphics representation of the simulated Fmoc-KYF hydrogels binding $O_2$. Characteristic binding modes of $O_2$ on the surface of Fmoc-KYF (binding mode I) and within the Fmoc-KYF nanofiber (binding mode II for the shallow layer and binding mode III for the deep layer) are indicated by solid black lines. (b) Interactions between the simulated Fmoc-KYF and $O_2$ in binding mode I. (c) Interactions between the simulated Fmoc-KYF hydrogels and $O_2$ in binding mode II. (d) Interactions between the simulated Fmoc-KYF hydrogels and $O_2$ in binding mode III. The Fmoc-KYF structures are visualized using licorice representation with transparent material, while interacting moieties are highlighted in CPK and licorice representation with opaque material. $O_2$ molecules are depicted in red color using van der Waals (VDW) representation.

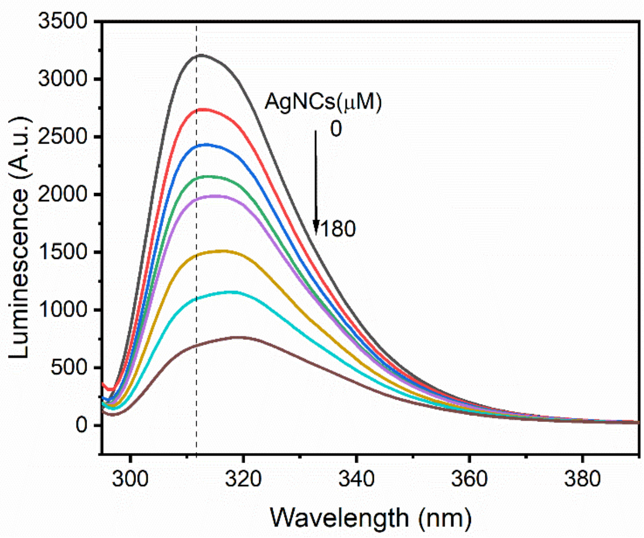


**Fig. S19**. Fluorescence spectra of the Fmoc-KYF peptide were recorded upon titration with AgNCs. The concentration of the Fmoc-KYF peptide was 5 mg mL$^{-1}$ , while the concentrations of AgNCs ranged from 0 to 180 μM. The excitation wavelength was set at 280 nm.

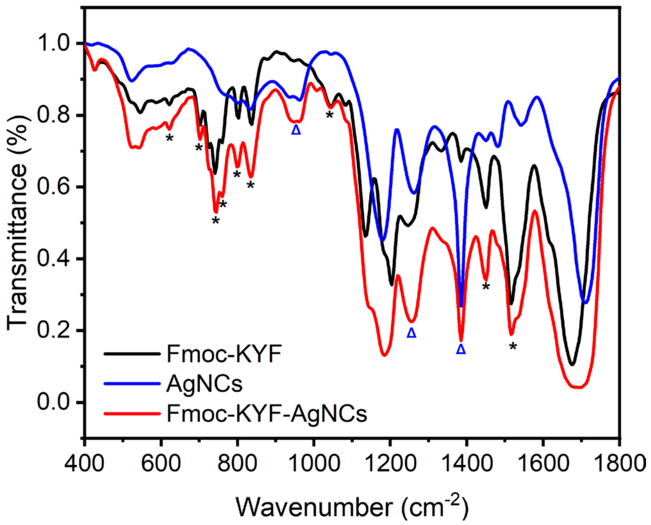


**Fig. S20**. FTIR spectra of AgNCs, Fmoc-KYF hydrogel and Fmoc-KYF- AgNCs hydrogel. The concentration of AgNCs is 0.5 mg mL$^{-1}$, concentrations of Fmoc-KYF is 5 mg mL$^{-1}$.

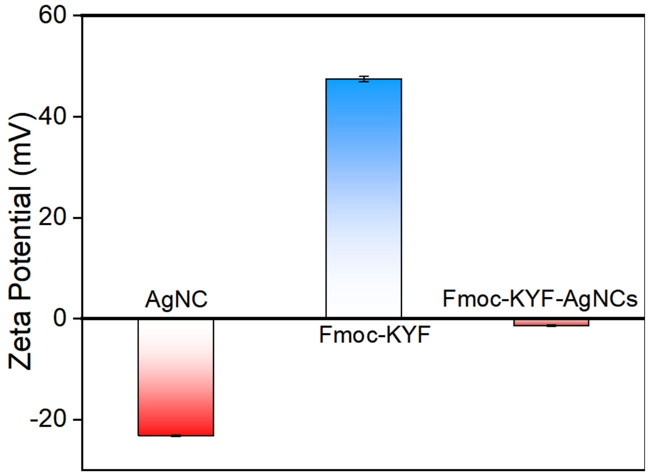


**Fig. S21.** ζ-potential measurements AgNCs, Fmoc-KYF hydrogel and Fmoc-KYF- AgNCs hydrogel. The concentration of AgNCs is 0.5 mg $mL^{-1}$ and Fmoc-KYF is 5 mg $mL^{-1}$.

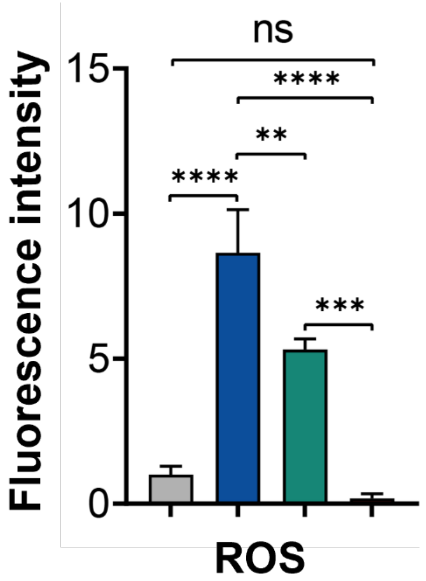


**Fig. S22.** Quantitative analysis of ROS levels in different groups. The control group is represented in gray, the LPS-treated group in blue, the LPS+Fmoc-KYF-AgNCs hydrogel-treated group in green, and the LPS+Fmoc-KYF-AgNCs-3S1C-Hyd-2 assembly-treated group in red. Statistical significance: ns, no significance, **P<0.01, ***P<0.001, ****P<0.0001, n=3 biologically independent samples.

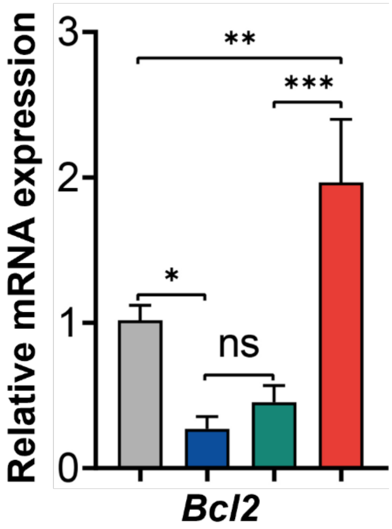


**Fig. S23.** Quantitative analysis of *anti-apoptotic* gene *Bcl2* levels in different groups. The control group is represented in gray, the LPS-treated group in blue, the LPS+Fmoc-KYF-AgNCs hydrogel-treated group in green, and the LPS+Fmoc-KYF-AgNCs-3S1C-Hyd-2 assembly-treated group in red. Statistical significance: ns, no significance, **P<0.01, ***P<0.001, ****P<0.0001, n=3 biologically independent samples.

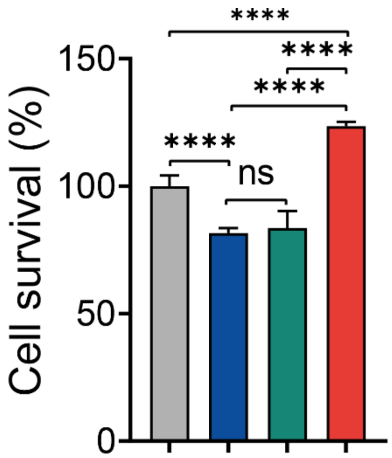


**Fig. S24.** CCK-8 assay of the cell viability. The control group is represented by gray, the LPS-treated group by blue, the LPS+Fmoc-KYF-AgNCs hydrogel-treated group by green, and the LPS+Fmoc-KYF-AgNCs-3S1C-Hyd-2 assembly-treated group by red. Statistical significance: ****P<0.0001.

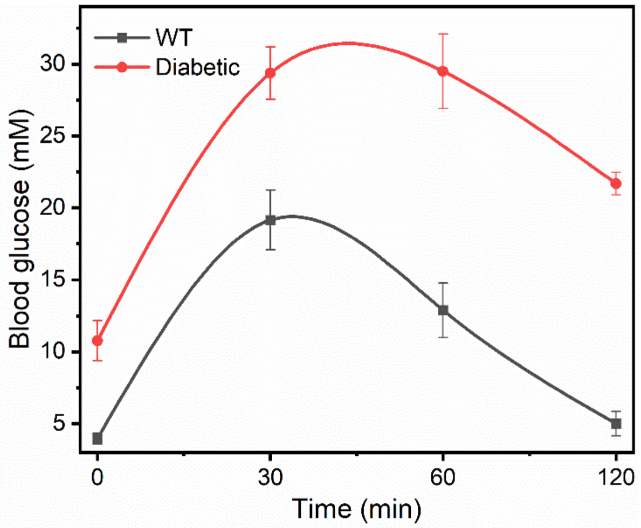


**Fig. S25.** Oral Glucose Tolerance Test. Following intragastric administration in mice, blood glucose concentrations were measured at 30-, 60-, and 120-minute intervals post-administration.

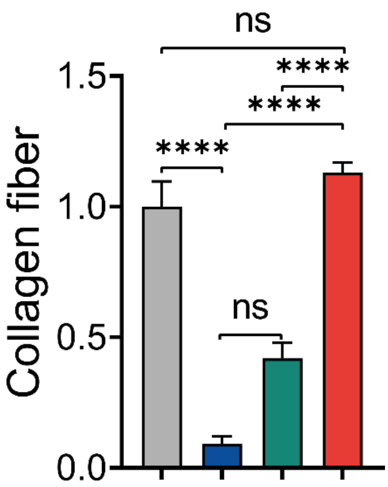


**Fig. S26**. Collagen deposition assay. The healthy mice group is represented by the gray color, the diabetic mice group by the blue color. The green color corresponds to the diabetic mice treated with Fmoc-KYF-AgNCs, while the red color represents the diabetic mice treated with Fmoc-KYF-AgNCs-3S1C-Hyd-2. Statistical significance: ns, no significance, ****P<0.0001.

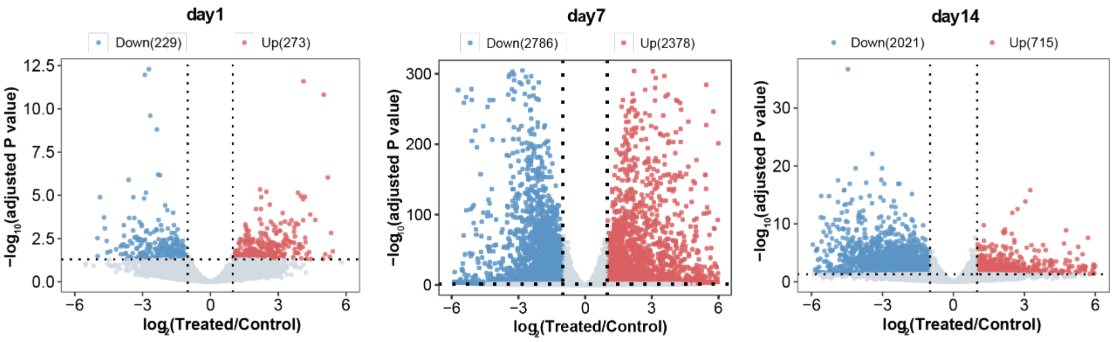


**Fig. S27**. Volcano plots illustrating the expression changes of differentially expressed genes (DEGs) between the treated and control groups on days 1, 7, and 14.

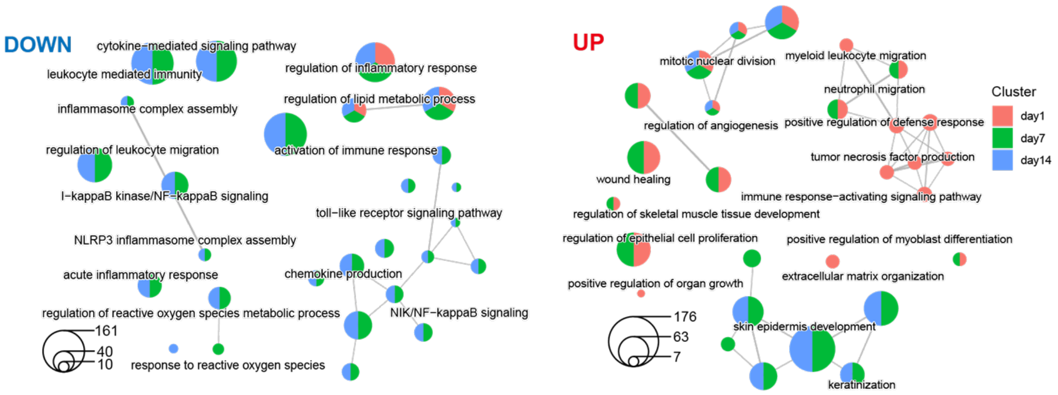


**Fig. S28**. Network visualization of representative Gene Ontology (GO) terms and pathways enriched in downregulated (left) and upregulated (right) genes between the treated and control groups on days 1, 7, and 14. Nodes are depicted as pie charts, with the size of each pie proportional to the total number of hits associated with the specific term. The pie charts are color-coded by pathways, where the size of each slice reflects the percentage of genes under the term that originate from the corresponding gene list.

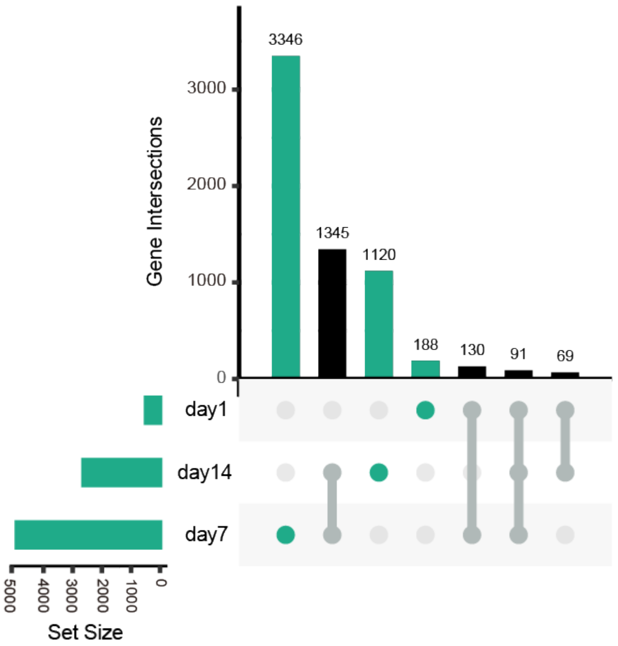


**Fig. S29**. UpSet plot showing the overlaps of differentially expressed genes (DEGs) across three time points (Day 1, Day 7, and Day 14).

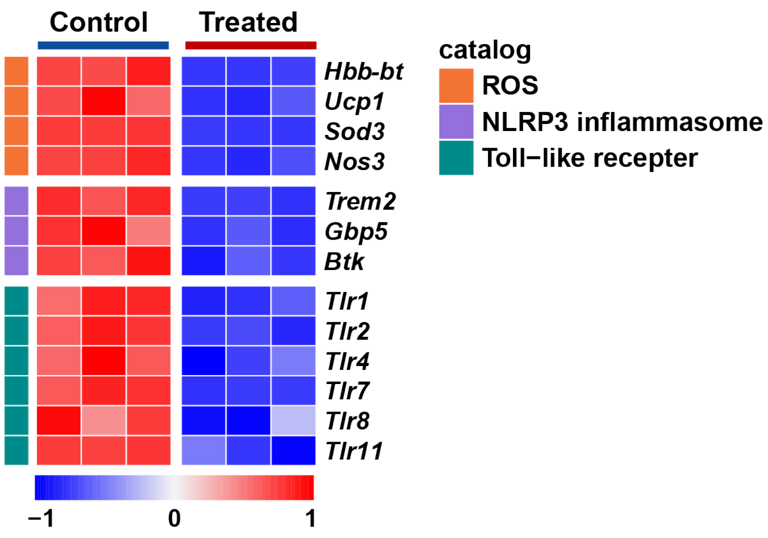


**Fig. S30** Heatmap illustrating the representative down-regulated expression of genes related to ROS, NLRP3 inflammasome, and Toll-like receptors in the treated group compared to the control group at Day 7.

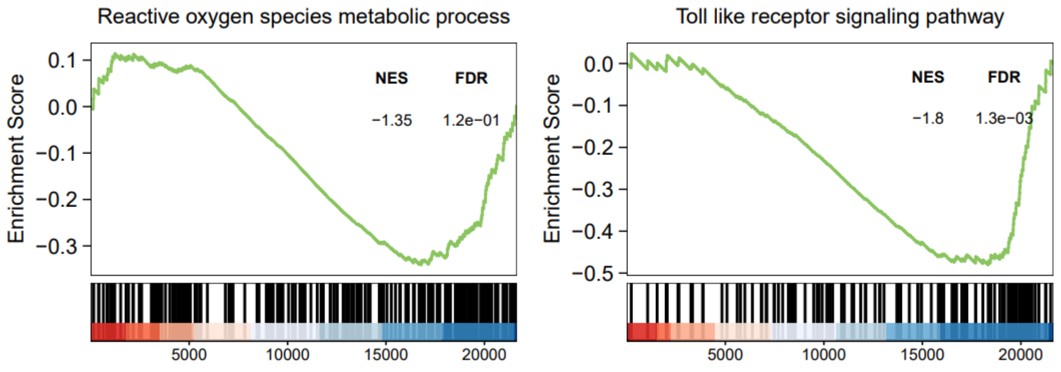


**Fig. S31**. GSEA plots depicting the ROS metabolic process pathway (left) and the toll-like receptor signaling pathway (right), comparing the treated group with the control group at Day 7. The normalized enrichment score (NES) and false discovery rate (FDR) are indicated.

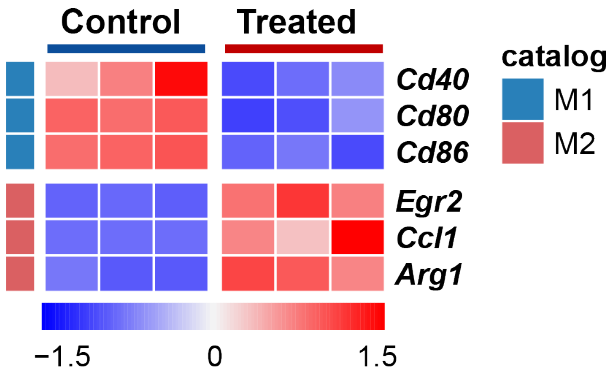


**Fig. S32**. Heatmap illustrating the gene expression levels of representative marker genes for M1 and M2 macrophages in both control and treated samples on day 7.

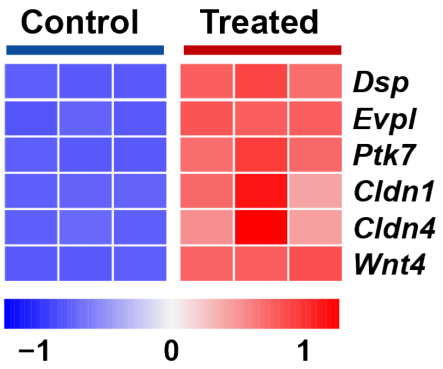


**Fig. S33**. Heatmap illustrating the up-regulated expression of representative wound-healing-associated genes in the treated group on day 7.

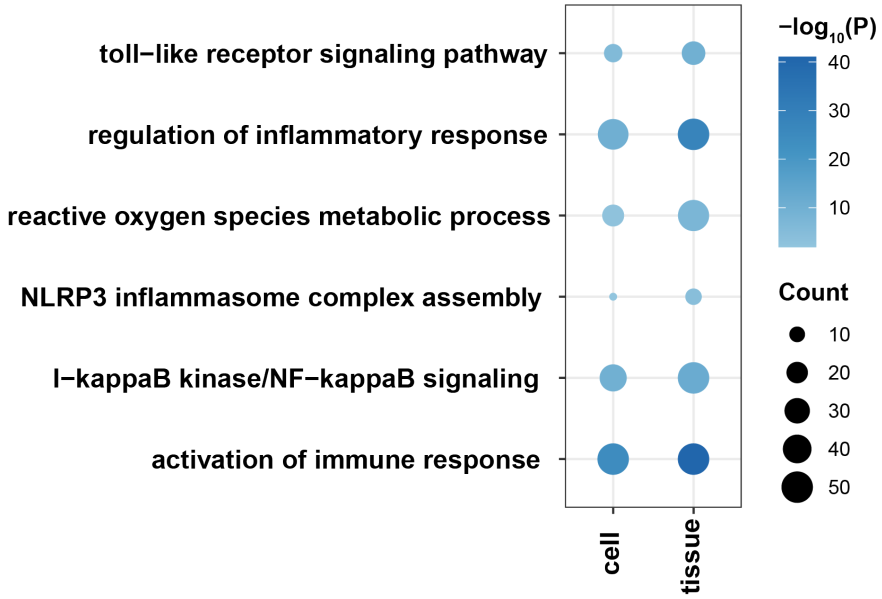


**Fig. S34**. Dot plot illustrating representative GO terms enriched in down-regulated genes in tissues and cells, comparing the treated to the control group on day 7.

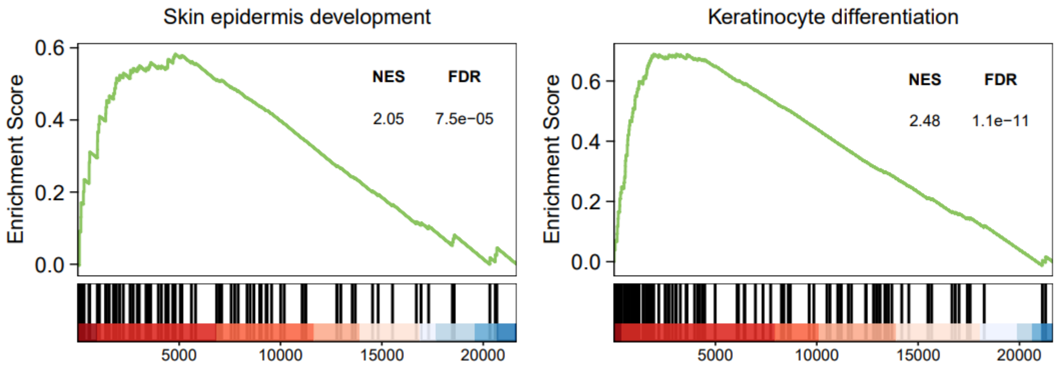


**Fig. S35**. GSEA plots depicting the "Skin epidermis development" (left) and "Keratinocyte differentiation" (right) pathways, comparing the treated group with the control group at Day 7. The NES and FDR are indicated.
